# Development of a mini-replicon-based reverse-genetics system for rice stripe tenuivirus

**DOI:** 10.1101/2021.04.04.438373

**Authors:** Mingfeng Feng, Luyao Li, Ruixiang Cheng, Yulong Yuan, Yongxin Dong, Minglong Chen, Rong Guo, Min Yao, Yi Xu, Yijun Zhou, Jianxiang Wu, Xin Shun Ding, Xueping Zhou, Xiaorong Tao

## Abstract

Negative-stranded RNA (NSR) viruses include both animal- and plant-infecting viruses that often cause serious diseases in human and livestock, and in agronomic crops. Rice stripe tenuivirus (RSV), a plant NSR virus with four negative-stranded/ambisense RNA segments, is one of the most destructive rice pathogens in many Asian countries. Due to the lack of a reliable reverse-genetics technology, molecular studies of RSV gene functions and its interaction with host plants are severely hampered. To overcome this obstacle, we developed a mini-replicon-based reverse-genetics system for RSV gene functional analysis in *Nicotiana benthamiana*. We first developed a mini-replicon system expressing RSV genomic RNA3 eGFP reporter (MR3_(-)eGFP_), a nucleocapsid (NP), and a codon usage optimized RNA-dependent RNA polymerase (RdRp_opt_), respectively. Using this mini-replicon system we determined that RSV NP and RdRp_opt_ are indispensable for the eGFP expression from MR3_(-)eGFP_. The expression of eGFP from MR3_(-)eGFP_ can be significantly enhanced in the presence of NSs and P19-HcPro-γb. In addition, NSvc4, the movement protein of RSV, facilitated eGFP trafficking between cells. We also developed an antigenomic RNA3-based replicon in *N. benthamiana.* However, we found that the RSV *NS3* coding sequence acts as a *cis*-element to regulate viral RNA expression. Finally, we made mini-replicons representing all four RSV genomic RNAs. This is the first mini-replicon-based reverse-genetics system for monocot-infecting tenuivirus. We believe that this mini-replicon system described here will allow the studies of RSV replication, transcription, cell-to-cell movement and host machinery underpinning RSV infection in plants.

**IMPORTANCE:** Plant-infecting segmented negative-stranded RNA (NSR) viruses are grouped into 3 genera: *Orthotospovirus, Tenuivirus* and *Emaravirus*. The reverse-genetics systems have been established for members in the genera *Orthotospovirus* and *Emaravirus*, respectively. However, there is still no reverse-genetics system available for *Tenuivirus*. Rice stripe virus (RSV) is a monocot-infecting tenuivirus with four negative-stranded/ambisense RNA segments. It is one of the most destructive rice pathogens and causes significant damages to rice industry in Asian countries. Due to the lack of a reliable reverse-genetics system, molecular characterizations of RSV gene functions and the host machinery underpinning RSV infection in plants are extremely difficult. To overcome this obstacle, we developed a mini-replicon-based reverse-genetics system for RSV in *Nicotiana benthamiana*. This is the first mini-replicon-based reverse-genetics system for tenuivirus. We consider that this system will provide researchers a new working platform to elucidate the molecular mechanisms dictating segmented tenuivirus infections in plant.

## Introduction

Negative-sense RNA (NSR) viruses include well-known members of medical importance such as Ebola virus (EBOV), vesicular stomatitis virus (VSV), influenza A virus (FLUAV) and Rift Valley fever virus (RVFV) (1, 2) and include serious plant pathogens of agrinomical importance such as Tomato spotted wilt virus (TSWV), Rice stripe virus (RSV), and Rose rosette virus (RRV) (3–5). There are 3 genera of segmented NSR viruses infecting plants: *Orthotospovirus, Tenuivirus* and *Emaravirus*. TSWV and RSV are the representative viruses for *Orthotospovirus and Tenuivirus*, respectively (6, 7). RRV and *European mountain ash ringspot-associated virus* (EMAraV) are important members in the genus *Emaravirus* (8, 9).

Tenuiviruses are classified in the genus *Tenuivirus*, family *Phenuiviridae* within the order *Bunyavirales*. Most viruses in the family *Phenuiviridae* infect animals. RSV is one of most devastating causal agents of rice and often causes severe damages to rice production in China and many other Asian countries (6, 10, 11). RSV is transmitted by small brown planthopper (*Laodelphax striatellus*) in a persistent and circulative-propagative manner (6, 12–14). RSV genome consists of four RNA segments and encodes seven proteins through an antisense or an ambisense coding strategy (15–18). RSV RNA1 is of negative polarity and encodes the RNA-dependent RNA polymerase (RdRp) (17). RSV RNA2 encodes the NS2 protein in the viral (v) strand and the NSvc2 protein from the viral complementary (vc) strand (19). The NS2 protein is a weak viral suppressor of RNA silencing (VSR) and is required for RSV systemic infection in plants. The NSvc2 protein is a putative glycoprotein that targets endoplasmic reticulum (ER) and Golgi apparatus, and function as a helper factor to conquer insect midgut barriers after being cleaved into two mature proteins (i.e., NSvc2-N, the amino-terminal half, and NSvc2-C, the carboxyl-terminal half) (12, 20). RSV RNA3 encodes the NS3 protein in the v strand, and is known as a VSR that binds single- and double-stranded RNAs to suppress RNA silencing (21, 22), and the NP protein in the vc strand that interacts with viral genomic RNAs to form viral ribonucleoprotein complexes (RNPs) (23). RSV RNA4 encodes the SP protein in the v strand, a nonstructural and disease-specific protein that interacts with a host oxygen-evolving complex protein to interfere host photosynthesis, and the NSvc4 protein in the vc strand, a protein involved in RSV cell-to-cell and long-distance movement in plant (4, 21). The four RSV genomic RNAs all contain highly conserved 5′- and 3′-untranslated regions (UTR), important for the initiation of viral RNA transcriptions. RSV RNA2, 3 and 4 all have a noncoding intergenic region (IGR) with multiple AU-rich regions that form secondary hairpin-like structures to act as transcription termination signals (24, 25).

Viral reverse-genetics systems are important tools for the studies of viral gene functions, disease inductions and host factors involved in virus infection in plants (26–32). Although reverse-genetics systems have been firstly reported for animal segmented negative-stranded RNA viruses over 20 years ago (26, 33–43), to establish similar systems for plant segmented negative-stranded/ambisense RNA viruses turned out to be very challenging. However, just recently the reverse-genetics systems have been established for a few of non-segmented and segmented plant NSR viruses. The first reverse-genetics system of the non-segmented plant NSR viruses was established for *Sonchus yellow net virus* (SYNV, a nucleorhabdovirus) and followed with BYSMV (a cytorhabdovirus) (27, 44–47). Recently we established the first reverse-genetics system of segmented plant NSR viruses for TSWV (48). Soon after, the reverse-genetics system for RRV was also established, allowing for studies of emaravirus gene function and disease pathology in whole plants (9).

For 3 genera of segmented NSR viruses infecting plants, reverse-genetics model has been established for 2 of the genera, *Orthotospovirus* and *Emaravirus* (9, 48, 49). However, there is still no reverse-genetics system available for the genus *Tenuivirus*. The recent progresses on TSWV and RRV reverse-genetics system encouraged us to establish a reverse-genetics system for RSV. In this study, we developed a mini-replicon-based reverse-genetics system for RSV gene function analyses in *Nicotiana benthamiana*. This represents the first mini-replicon-based reverse-genetics system for monocot-infecting tenuivirus. The developed mini-replicon reverse-genetics system will provide researchers a novel platform for studies of RSV replication, transcription, movement and host factors involved in the interactions between the virus and the host plant. This system also provides useful basis for the development of infectious RSV clones for the assays in plants, including rice.

## Results

### Development of a genomic RNA3 mini-replicon-based reverse-genetics system for RSV

To establish a mini-replicon-based reverse-genetics system to investigate RSV infection in plant, we RT-PCR-amplified the full-length RSV RNA3 sequence and inserted it between an HH sequence and an RZ sequence in the pCB301-2×35S-RZ-NOS vector to produce RNA3_(-)_. We then replaced the *NP* gene in RNA3_(-)_ with an *eGFP* gene to produce MR3_(-)eGFP_ mini-replicon reporter (Fig. 1A). The expressions of RNA3_(-)_ and MR3_(-)eGFP_ from these two vectors are driven by a doubled *Cauliflower mosaic virus* (CaMV) 35S promoter (2×35S) (Fig. 1A).

**Fig. 1.**
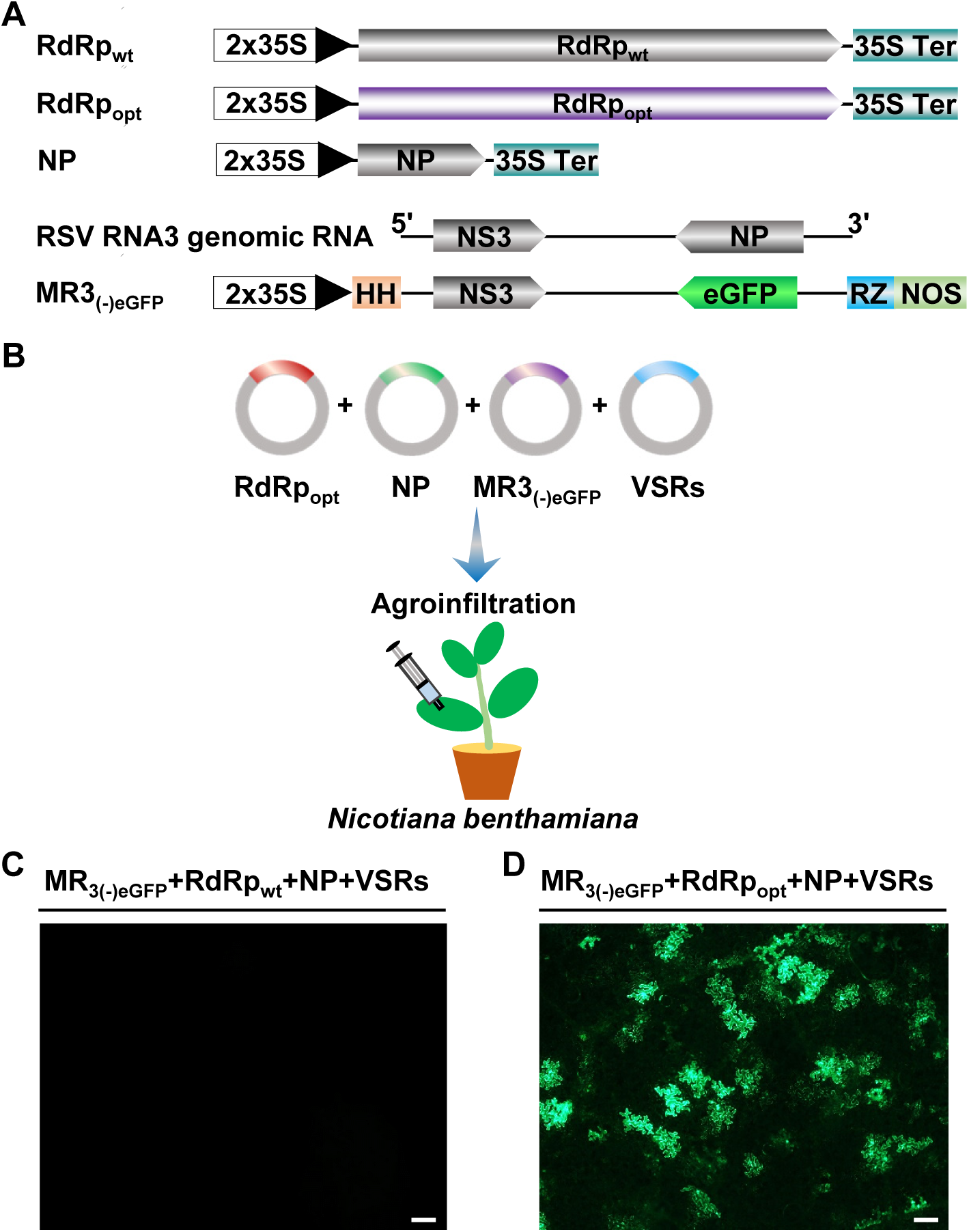
Construction of an RSV RNA3_(-)_-based mini-replicon. (A) Schematics representing the RdRp_wt_, RdRp_opt_, NP, RSV RNA3_(-)_ and MR3_(-)eGFP_ mini-replicong. For MR3_(-)eGFP_, we replaced the *NP* gene in the gRNA3 with an *eGFP* gene. The 5′ untranslated, the 3′ untranslated, and the intergenic region in RNA3_(-)_ were indicated with a thin black line. 2×35S, doubled 35S promoter; HH, hammerhead ribozyme; RZ, hepatitis delta virus (HDV) ribozyme; 35S Ter, 35S terminator; NOS, nopaline synthase terminator. Minus sign (-) and 5′ to 3′ designation represent the viral (genomic)-strand of RNA3. (B) Illustration of agro-infiltration using a mixed Agrobacterium culture carrying various min-replicons into *N. benthamiana* leaves. VSRs, NSs plus P19-HcPro-γb. (C) A *N. benthamiana* leaf infiltrated with a mixed Agrobacterium culture carrying MR3_(-)eGFP_, RdRp_wt_, NP and VSRs (NSs and P19-HcPro-γb). (D) A *N. benthamiana* leaf infiltrated with a mixed Agrobacterium culture carrying MR3_(-)eGFP_, RdRp_opt_, NP and VSRs (NSs and P19-HcPro-γb) and was showing eGFP fluorescence at 5 dpi under an inverted fluorescence microscope. Bar represents 200 μm.

We constructed p2300-RdRp_wt_, pBIN-NS3, p2300-NP, and pCXSN-NSvc4 to express the wild-type RSV RdRp (RdRp_wt_), NS3, NP, and NSvc4, respectively, in cells through agro-infiltration. After infiltration of a mixed Agrobacterium culture carrying the plasmids of MR3_(-)eGFP_, RdRp_wt_, NP and four VSRs (NSs and P19-HcPro-γb) into *N. benthamiana* leaves (Fig 1B), no eGEP fluorescence was observed in the infiltrated leaves by 5 days post agro-infiltration (dpi) (Fig. 1C). As reported for TSWV RdRp (48), the wild-type RSV RdRp was also predicted to have numerous putative intron splicing sites. This prediction prompted us to optimize its codon usage and to remove the predicted intron splicing sites. The optimized *RdRp* ORF was then inserted into the p2300 vector to produce p2300-RdRp_opt_ (RdRp_opt_). After infiltrating *N. benthamiana* leaves with a mixed Agrobacterium culture carrying plasmids of MR3_(-)eGFP_, RdRp_opt_, NP and four VSRs, we observed the eGFP fluorescence using a microscope at 5 dpi (Fig. 1B), suggesting that the sequence optimized RdRp_opt_ is now functional in *N. benthamiana* cells.

### RSV NP and RdRp_opt_ are required for MR3_(-)eGFP_ expression

To investigate the roles of NP and RdRp_opt_ in RSV replication in plant, we co-expressed MR3_(-)eGFP_ and VSRs with empty vector (Vec), NP, RdRp_opt_, or both NP and RdRp_opt_, respectively, in *N. benthamiana* leaves through agro-infiltration. The eGFP fluorescence was again detected in the leaves co-expressing MR3_(-)eGFP_ with NP and RdRp_opt_. In contrast, the control leaves expressing MR3_(-)eGFP_ alone, or co-expressing MR3_(-)eGFP_ with NP or RdRp_opt_, did not show any eGFP fluorescence (Fig. 2A), indicating that the presence of both NP and RdRp_opt_ is necessary for MR3_(-)eGFP_ expression from the mini-replicon.

**Fig. 2.**
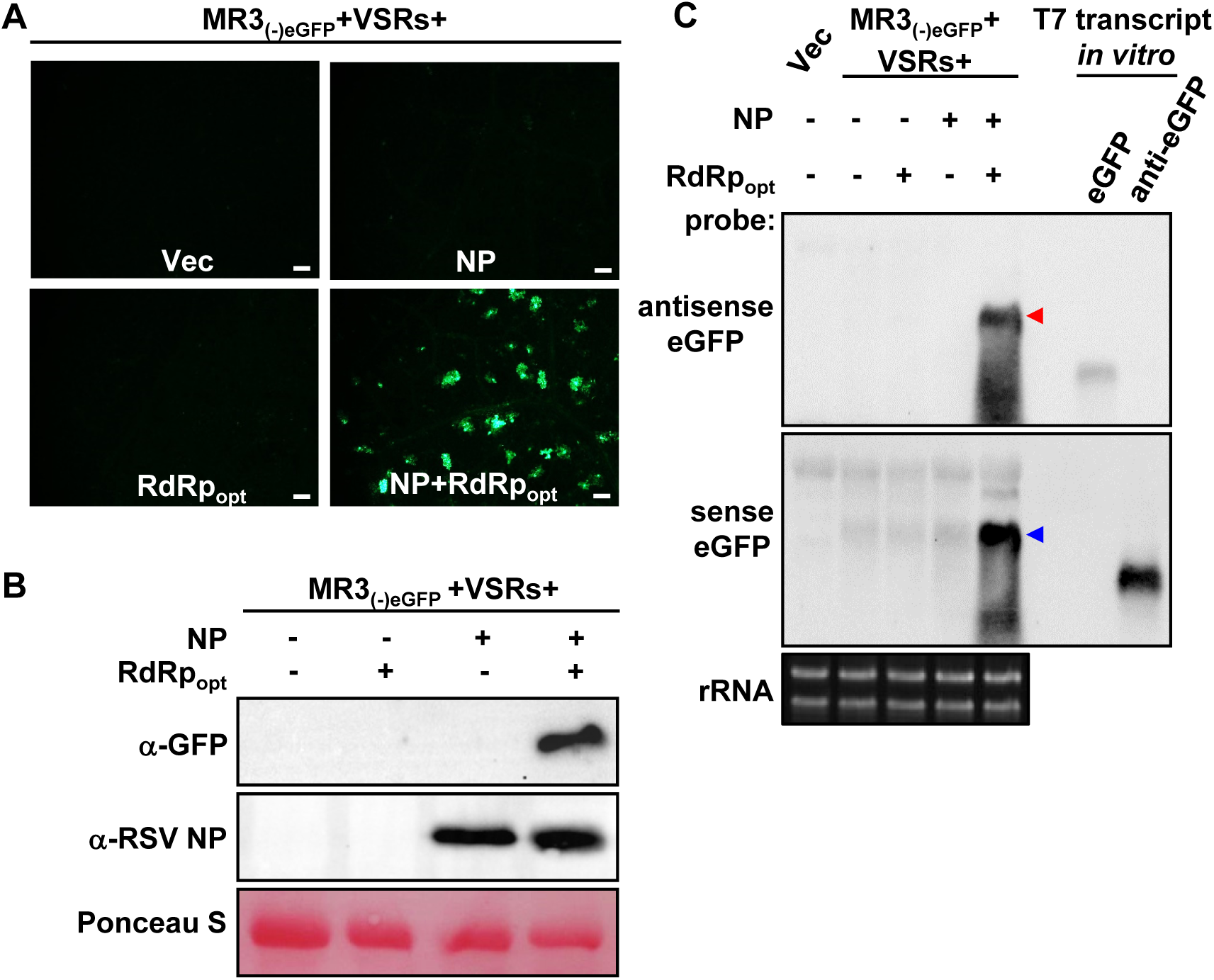
RSV NP and RdRp_opt_ are required for MR3_(-)eGFP_ expression in *N. benthamiana* leaves. (A) *N. benthamiana* leaves were infiltrated with mixed *Agrobacterium* cultures carrying MR3_(-)eGFP_+NSs+P19-HcPro-γb+Vec (empty vector), MR3_(-)eGFP_+NSs+P19-HcPro-γb+NP, MR3_(-)eGFP_+NSs+P19-HcPro-γb+RdRp_opt_, or MR3_(-)eGFP_+NSs+P19-HcPro-γb+RdRp_opt_. The infiltrated leaves were examined and photographed under an inverted fluorescence microscope at 5 dpi. Bars = 200 μm. (B) Western blot analyses using the samples described in (A) and an NP or an eGFP specific antibody. The ponceau S-stained Rubisco large subunit gel was used to show sample loadings. (C) Northern blot analyses using the samples described in (A) and a DIG-labeled sense- or an antisense-eGFP probe. The red and blue arrows indicate the antigenomic and genomic RNA3 expressed the infiltrated leaves. The ethidium bromide stained ribosomal RNA gel was used to show sample loadings.

To confirm this observation, we analyzed the accumulation levels of eGFP protein, *eGFP* mRNA, and MR3_(-)eGFP_ genomic RNA (gRNA) and anti-genomic RNA (agRNA) in the infiltrated *N. benthamiana* leaf tissues. Western blot showed high levels of eGFP in the leaves co-expressing MR3_(-)eGFP_, NP, RdRp_opt_ and four VSRs (NSs+P19-HcPro-γb). In contrast, eGFP protein was not detected in the control leaves co-expressing MR3_(-)eGFP_ with Vec, NP or RdRp_opt_ only by 5 dpi (Fig. 2B). Northern blot results showed that the gRNA and agRNA were detected in the leaf tissues co-expressing MR3_(-)eGFP_, NP, RdRp_opt_ and VSRs by 5 dpi (Fig. 2C). In contrast, no amplified gRNA and agRNA were detected in the control leaves co-expressing MR3_(-)eGFP_ with Vec, NP or RdRp only, respectively (Fig. 2C). Only primary transcripts of agRNA from MR3_(-)eGFP_ was detected in these leaves. These results indicate that the presence of RSV NP and RdRp_opt_ are required for the replication of both gRNA and agRNA from the MR3_(-)eGFP_ mini-replicon. Intrinsically, the *eGFP* mRNA was not detected in the leaf tissues co-expressing MR3_(-)eGFP_, NP, RdRp_opt_ and VSRs.

### Effect of viral suppressors of RNA silencing on MR3_(-)eGFP_ expression

Viral suppressors of RNA silencing (VSRs) inhibit host RNAi machinery and enhance non-viral gene expressions in plants (44, 46). To investigate the roles of different VSRs on MR3_(-)eGFP_ expression in plant, we infiltrated *N. benthamiana* leaves with mixed agrobacterium cultures carrying MR3_(-)eGFP_, RdRp_opt_, NP and VSRs including NS3, NSs, and P19-HcPro-γb, respectively. By 5 dpi, the leaves co-expressing MR3_(-)eGFP_, RdRp_opt_ and NP with Vec showed a few cells with eGFP fluorescence. Many cells with eGFP fluorescence were observed in the leaves co-expressing MR3_(-)eGFP_, RdRp_opt_ and NP with three VSRs (P19-HcPro-γb) or four VSRs (NSs and P19-HcPro-γb) (Fig. 3A and 3B). Although RSV NS3 is a VSR (21, 22), co-expression of MR3_(-)eGFP_, RdRp_opt_, NP with NS3 or both NS3 and NSs resulted in almost no cell with eGFP fluorescence. Moreover, co-expression of MR3_(-)eGFP_, RdRp_opt_, NP with both NS3 and P19-HcPro-γb resulted in some cells with eGFP fluorescence. These findings indicate that RSV NS3 can suppress eGFP expression from the MR3_(-)eGFP_ mini-replicon, and are supported by the Western blot result (Fig. 3B). Based on these results, we decided to use VSRs including NSs, P19, HcPro and γb but not NS3 in the following experiments.

**Fig. 3.**
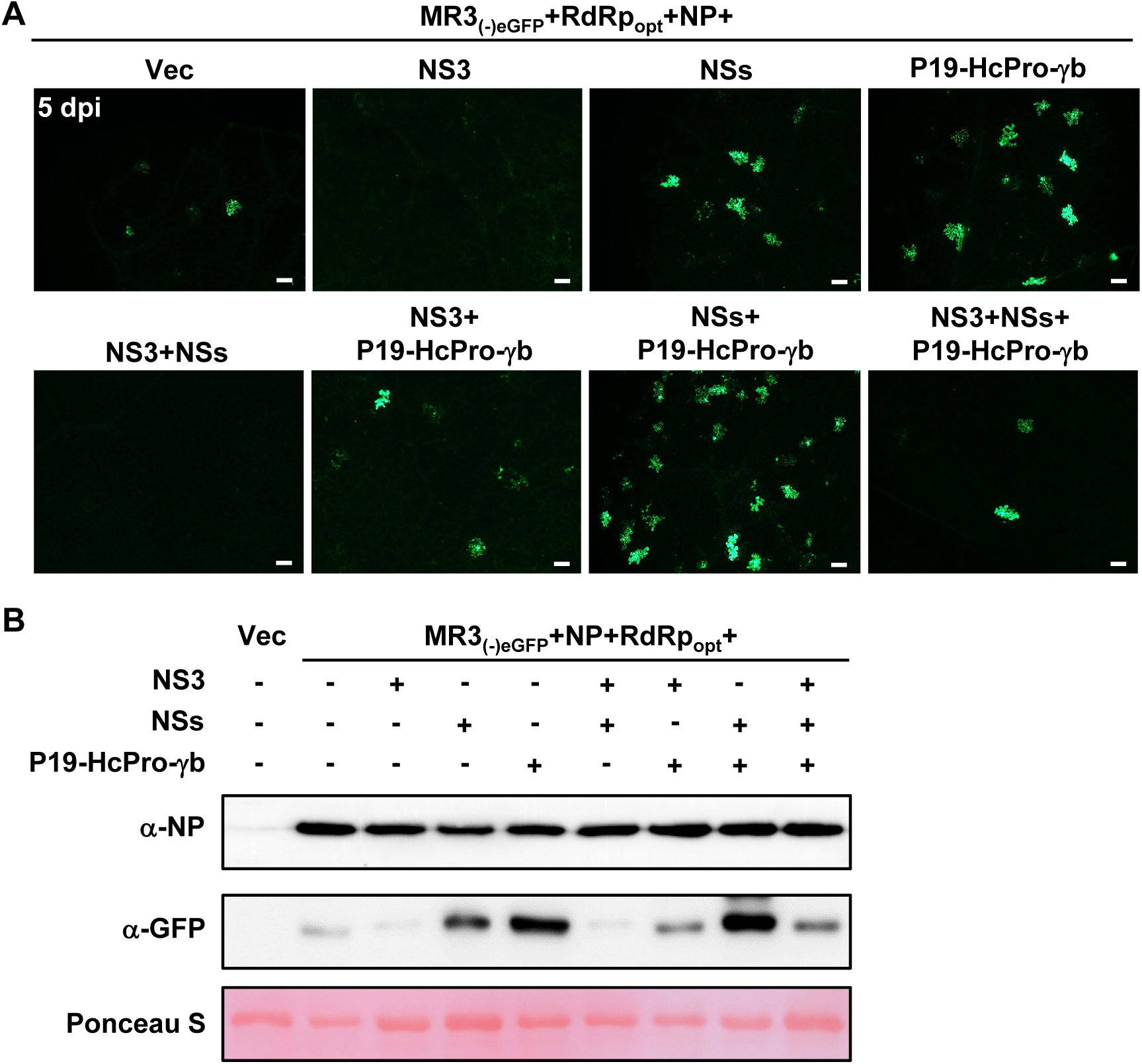
Effects of VSRs on pMR3_(-)eGFP_ expression. (A) *N. benthamiana* leaves were infiltrated with mixed Agrobacterium cultures as indicated in the figure. The infiltrated *N. benthamiana* leaves were harvested at 5 dpi, and examined and photographed under an inverted fluorescence microscope. Bars = 200 μ Western blot analyses using the samples described in (A), and an NP specific and an eGFP specific antibodies, respectively. Proteins in the leaves shown in panel (A) using NP and GFP-specific antibodies, respectively. The ponceau S-stained Rubisco large subunit gel was used to show sample loadings.

### Dosage effects of NP and RdRp_opt_ on MR3_(-)eGFP_expression

To optimize the expression of MR3_(-)eGFP_ mini-replicon in plant cells, we mixed the Agrobacterium culture carrying MR3_(-)eGFP_ and four VSRs (NSs+P19-HcPro-γb) with the Agrobacterium cultures carrying NP (OD_600_ 0, 0.05, 0.1, 0.2 or 0.4) or RdRp_opt_ (OD_600_ 0, 0.05, 0.1, 0.2 or 0.4), and then infiltrated them individually into *N. benthamiana* leaves. When the concentration of Agrobacterium culture carrying RdRp was fixed at OD_600_ 0.05 and the concentration of Agrobacterium culture carrying NP was increased from OD_600_ 0.05 to OD_600_ 0.4, the results showed that the strongest eGFP fluorescence was observed in the leaves co-expressing MR3_(-)eGFP_ with NP at OD_600_ 0.2 and RdRp_opt_ at OD_600_ 0.05 (Fig. 4A). When the concentration of Agrobacterium culture carrying NP was maintained at OD_600_ 0.2, while the concentration of Agrobacterium culture carrying RdRp_opt_ was increased from OD_600_ 0.05 to OD_600_ 0.4, the number of cells with eGFP fluorescence decreased as the concentration of RdRp_opt_ increased (Fig. 4B). These results were supported by the results from Western blot assays (Fig. 4C and 4D).

**Fig. 4.**
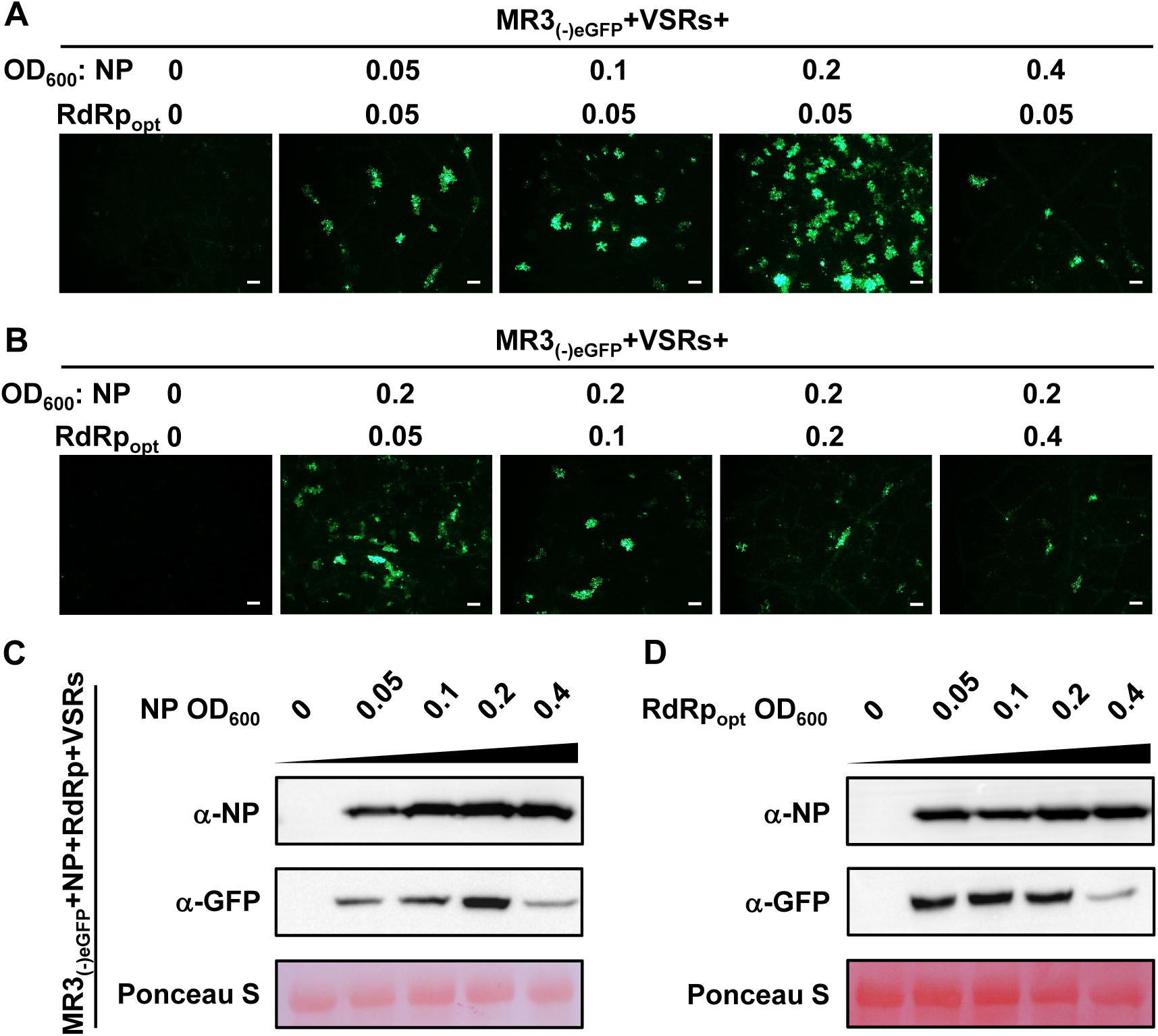
Concentrations of NP and RdRp_opt_ needed for the maximum expression MR3_(-)eGFP_. (A and B) *N. benthamiana* leaves were infiltrated with various mixed Agrobacterium cultures as indicated in the figure. The concentration of Agrobacterium cultures carrying NP or RdRp_opt_ ranged from OD_600_ = 0 to 0.4. The infiltrated leaves were harvested at 5 dpi, and examined and photographed under an inverted fluorescence microscope. Bars = 200 μm. (C and D) Western blot analyses of NP and eGFP expressions in the infiltrated leaves described in (A and B) using an NP specific and an eGFP specific antibodies, respectively. The ponceau S-stained Rubisco large subunit gel was used to show sample loadings.

### RSV NSvc4 supports MR3_(-)eGFP_ cell-to-cell movement

NSvc4 is the movement protein of RSV (18, 50). To investigate whether NSvc4 can also influence MR3_(-)eGFP_ expression, we infiltrated *N. benthamiana* leaves with the mixed Agrobacterium cultures carrying MR3_(-)eGFP_ (OD_600_ 0.2), NP (OD_600_ 0.2), RdRp_opt_ (OD_600_ 0.05), NSs (OD_600_ 0.05), P19-HcPro-γb (OD_600_ 0.05) and NSvc4 (OD_600_ 0.025, 0.05, 0.1 or 0.15). In this experiment, the MR3_(-)eGFP_ started to move out the original cell in the addition of NSvc4 at OD_600_ 0.025, compared with that in the leaves co-expressing MR3_(-)eGFP_, NP and RdRp_opt_ without NSvc4 (Fig. 5A and Table 1). When the concentration of Agrobacterium culture carrying NSvc4 was further increased, stronger cell-to-cell movement of MR3_(-)eGFP_ was observed (Fig. 5A and Table 1). Results of the Western blot assays agreed with the microscopic observations (Fig. 5B).

**Fig. 5.**
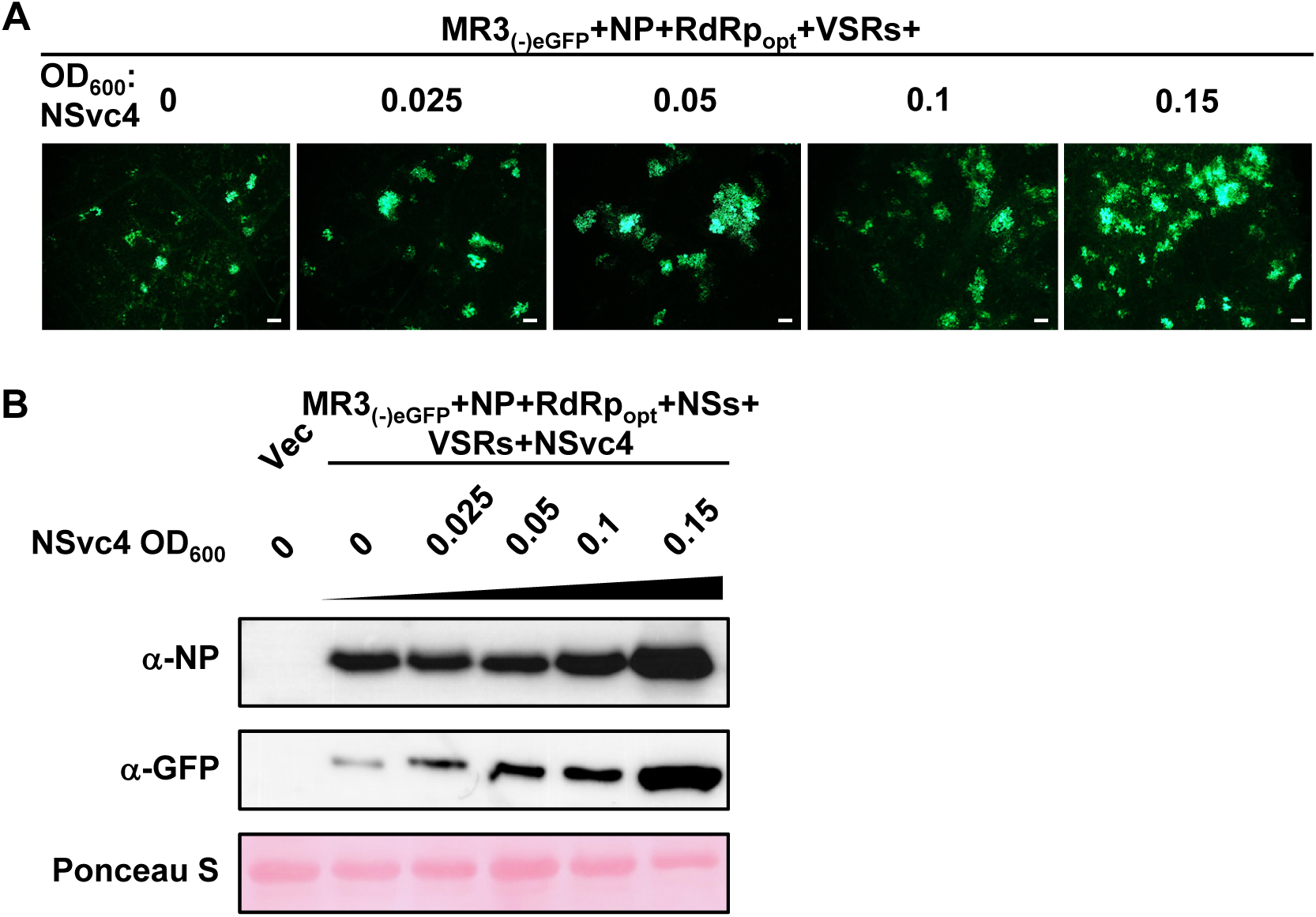
Effect of RSV NSvc4 on eGFP expression from MR3_(-)eGFP_ in cells. (A) *N. benthamiana* leaves were infiltrated with various mixed Agrobacterium cultures as described in the figure. The concentration of Agrobacterium culture carrying NSvc4 ranged from OD_600_ = 0 to 0.15. The infiltrated leaves were harvested at 5 dpi, and examined and photographed under an inverted fluorescence microscope. Bars = 200 μm. (B) Western blot analyses using the infiltrated leaf samples described in (A), and an NP specific and an eGFP specific antibodies, respectively. The ponceau S-stained Rubisco large subunit gel was used to show sample loadings.

**Table 1.** RSV NSvc4 enhances the cell-to-cell movement of MR3_(-)eGFP_ mini-replicon in the presence of NP, RdRp_opt_ and four VSRs.

| NSvc4<sup>a</sup><br>OD<sub>600</sub> | Total eGFP<br>fluorescent cells | No. of clusters with<br>1 cell (% of total) | No. of clusters with<br>2 cells (% of total) | No. of clusters with<br>3≥cells (% of total) |
| --- | --- | --- | --- | --- |
| 0 | 18 | 18 (100%)<sup>b</sup> | 0 (100) | 0 (100) |
| 0.025 | 54 | 15 (27.8%) | 6 (22.2%) | 5 (50%) |
| 0.05 | 76 | 12 (15.8%) | 7 (18.4%) | 10 (65.8%) |
| 0.1 | 95 | 8 (8.4%) | 11 (23.2%) | 13 (68.4%) |
| 0.15 | 140 | 17 (12.1%) | 9 (12.9%) | 18 (75%) |

<sup>a</sup> Agrobacterium cultures harboring the plasmid encoding NSvc4 (final concentration, OD<sub>600</sub> = 0, 0.025, 0.05, 0.1 and 0.15) were used to infiltrate *N. benthamiana* leaves.
<sup>b</sup> eGFP fluorescent foci of RSV MR3<sub>(-)eGFP</sub> at 5 days post-infiltration.

### Development of an RSV antigenomic RNA3-based replicon system

To develop an RSV antigenomic (ag) RNA-based mini-replicon, we replaced the *NS3* gene in the RNA3_(+)_-agRNA vector (RNA3_(+)_) with an *eGFP* gene to produce MR3_(+)eGFP_ (Fig. 6A). We then transiently co-expressed MR3_(+)eGFP_, NP, RdRp_opt_, and four VSRs (NSs+P19-HcPro-γb) in *N. benthamiana* leaves through agro-infiltration. By 5 dpi, no eGFP fluorescence was observed in the infiltrated leaves. In contrast, strong eGFP fluorescence was observed in the leaves co-expressing MR3_(-)eGFP_, NP, RdRp_opt_ and four VSRs (Fig. 6B), indicating that the MR3_(+)eGFP_ min-replicon is incapable of expressing eGFP in plant cells.

**Fig. 6.**
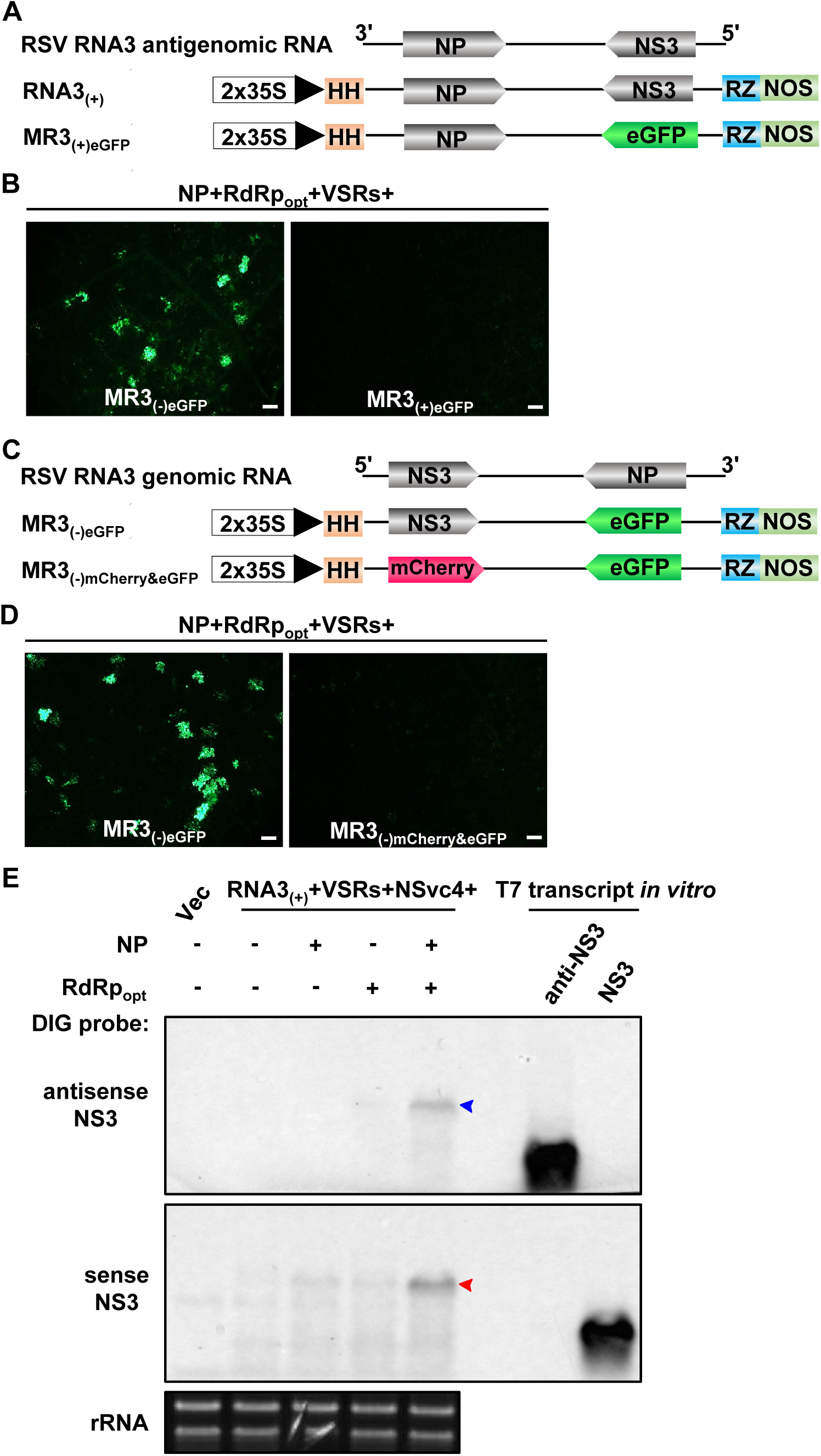
Construction and test of RNA3_(+)_-based mini-replicon in *N. benthamiana* leaves. (A) Schematics representing RNA3_(+)_ and MR3_(+)eGFP_ mini-replicons. The MR3_(+)eGFP_ mini-replicon was made by replacing the *NS3* gene in RNA3_(+)_ with an *eGFP* gene. Plus sign (+) and 3′ to 5′ designation represent the viral complementary (antigenomic)-strand of RNA3. (B) *N. benthamiana* leaves were infiltrated with various mixed Agrobacterium cultures as described in the figure. The infiltrated leaves were observed at 5 dpi, and examined and photographed under an inverted fluorescence microscope. Bars = 200 μm. (C) Schematics representing MR3_(-)eGFP_ and MR3_(-)mCherry&eGFP_ mini-replicons. The MR3_(-)mCherry&eGFP_ mini-replicon was constructed by replacing the *NS3* gene with an *mCherry* gene. Minus sign (-) and 5′ to 3′ designation represent the viral (genomic)-strand of RNA3. (D) The infiltrated *N. benthamiana* leaves were harvested at 5 dpi, and examined and photographed under an inverted fluorescence microscope. Bars = 200 μm. (E) Northern blot analyses of RNA3 expressions from RNA3_(+)_ in the infiltrated leaves as described in (C). After electrophoresis, the blots were probed with a DIG-labeled antisense and a sense RNA3 probes, respectively. The red and blue arrow heads indicate the RNA3 bands probed with DIG-labeled probes. The ethidium bromide stained ribosomal RNA (rRNA) gel was used to show sample loadings.

Next, we replaced the *NS3* gene in MR3_(-)eGFP_ with a *mCherry* gene (MR3_(-)mCherry&eGFP_) (Fig. 6C), and co-expressed MR3_(-)mCherry&eGFP_ or MR3_(-)eGFP_ with NP, RdRp_opt_ and four VSRs in *N. benthamiana* leaves through agro-infiltration. By 5 dpi, in contrast to the leaves co-expressing MR3_(-)eGFP_, NP, RdRp_opt_ and VSRs, no eGFP fluorescence was observed for MR3_(-)mCherry&eGFP_ (Fig. 6D), indicating that without the *NS3* gene, this MR3_(-)mCherry&eGFP_ min-replicon is unable to express eGFP in *N. benthamiana* leaf cells. To confirm this result, we generated RNA3_(+)_ to express full-length RSV agRNA3 (Fig. 6A), and transiently co-expressed RNA3_(+)_, VSRs, NSvc4 with one or two of the three plasmids (i.e., Vec, NP and RdRp_opt_), respectively in *N. benthamiana* leaves via agro-infiltration. The infiltrated leaves were harvested at 5 dpi and analyzed for the accumulations of RNA3_(+)_-derived gRNA3 and agRNA3 through Northern blot assays using DIG labelled antisense- and sense-probes, respectively. The result showed that high levels of gRNA3 and agRNA3 were detected in the leaves co-expressing RNA3_(+)_, NP, RdRp_opt_, VSRs and NSvc4 (Fig. 6E). In contrast, No amplified gRNA3 and agRNA3were detected in the leaves co-expressing RNA3_(+)_, four VSRs, NSvc4 with Vec, NP or RdRp_opt_ only) (Fig. 6E). Only primary transcripts of agRNA3 from RNA3_(+)_ was detected in these leaves. Based on these results, we conclude that the RNA3_(+)_ replicon is functional in *N. benthamiana* in the presence of NP, RdRp_opt_, NSs, P19-HcPro-γb and NSvc4.

### The *NS3* gene is required for *eGFP* expression from MR3_(-)eGFP_

To investigate the function of *NS3* in eGFP expression from the MR3_(-)eGFP_ mini-replicon, we introduced a stop codon (TAA) at the downstream of the start codon of *NS3* ORF (MR3_(-)eGFP&NS3stop_) (Fig. 7A), and co-expressed it with NP, RdRp_opt_ and VSRs in *N. benthamiana* leaves through agroinfiltration. By 5 dpi, strong eGFP fluorescence was observed in the infiltrated leaves, while the leaves co-expressing MR3_(-)eGFPΔNS3_, NP, RdRp_opt_ and VSRs did not (Fig. 7B). Western blot results showed that the eGFP were expressed in the leaves co-expressing MR3_(-)eGFP&NS3stop_, NP, RdRp_opt_ and VSRs, while no eGFP was accumulated in the leaves co-expressing MR3_(-)eGFPΔ NS3_, NP, RdRp_opt_ and VSRs (Fig. 7C). This finding indicates that the NS3 is dispensable for eGFP expression from the MR3_(-)eGFP_ mini-replicon. Deletion of *NS3* gene sequence from MR3_(-)eGFP_ (MR3_(-)eGFPΔNS3_) abolished the expression of eGFP.

**Fig. 7.**
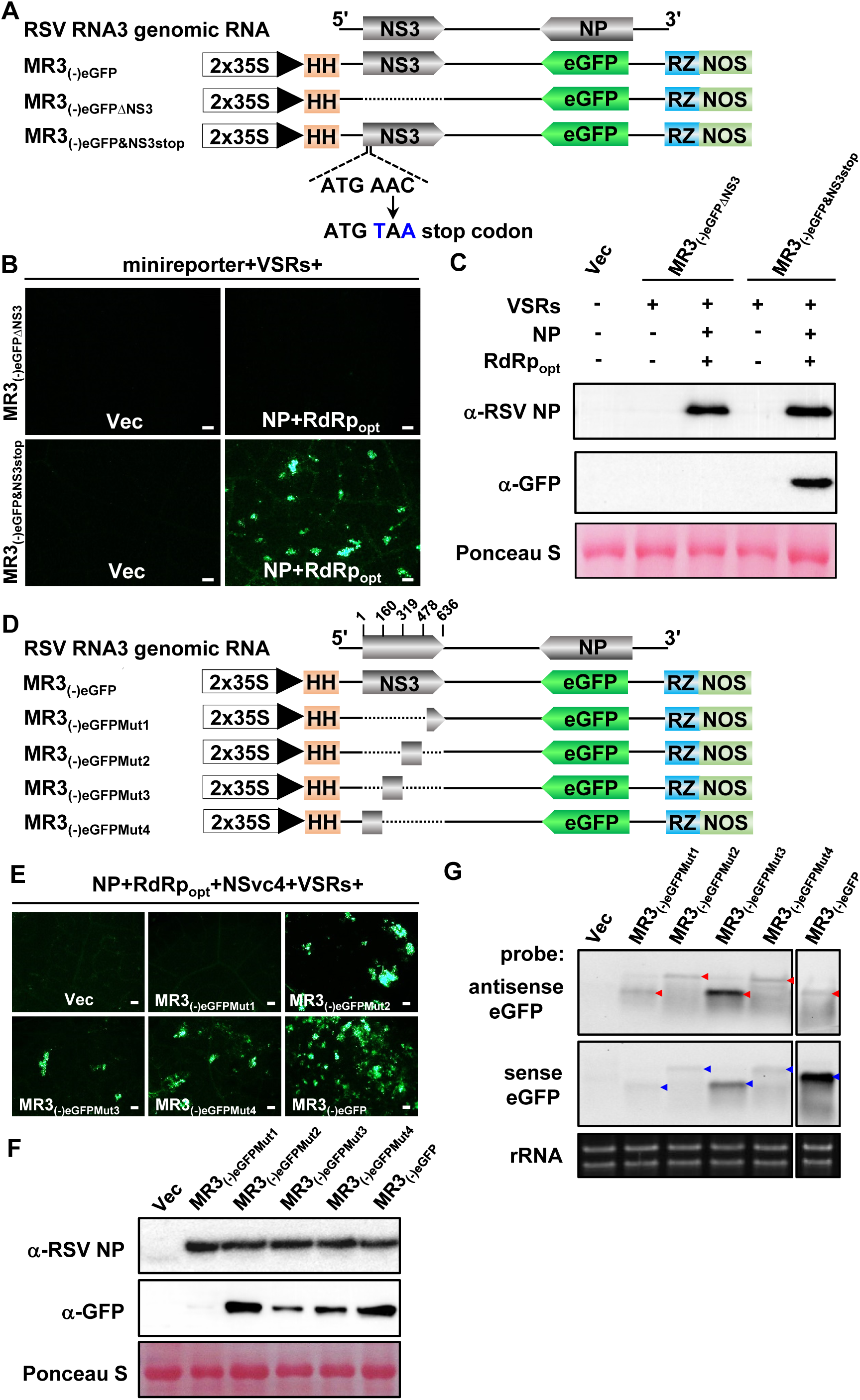
Effect of NS3 on *eGFP* gene expression from the mini-replicons. (A and D) Schematics representing MR3_(-)eGFP_ and its mutants. For MR3_(-)eGFPΔNS3_, the *NS3* gene was deleted from MR3_(-)eGFP_. For MR3_(-)eGFP&NS3stop_, a stop codon was introduced at the downstream of the start codon of the *NS3* gene in pMR3_(-)eGFP_. For MR3_(-)eGFPMut1 to 4_ mutant mini-replicons, a quarter of the *NS3* gene in MR3_(-)eGFP_ was deleted as shown in the figure. Minus sign (-) and 5′ to 3′ designation represent the viral (genomic)-strand of RNA3. (B and E) *N. benthamiana* leaves were infiltrated with various mixed Agrobacterium cultures as described in the figure. The infiltrated leaves were harvested at 5 dpi, and examined and photographed under an inverted fluorescence microscope. Bars = 200 μm. (C and F) Western blot analyses using the leaf samples described in (B and E), and an NP specific and an eGFP specific antibodies, respectively. The ponceau S-stained Rubisco large subunit gels were used to show sample loadings. (G) Northern blot analyses of antigenomic and genomic RNA expressions from MR3_(-)eGFP_ using a DIG-labeled antisense- and a sense-eGFP probe, respectively. The ethidium bromide-stained ribosomal RNA gel was used to show sample loadings.

To further confirm the role of *NS3* ORF in MR3_(-)eGFP_ expression, we divided *NS3* ORF into four segments and generated four truncated MR3_(-)eGFP_ mutant constructs: MR3_(-)eGFPMut1_, MR3_(-)eGFPMut2_, MR3_(-)eGFPMut3_, and MR3_(-)eGFPMut4_ (Fig. 7D). Each mutant was co-expressed with NP, RdRp_opt_, NSvc4 and VSRs, respectively, in *N. benthamiana* leaves. By 5 dpi, eGFP fluorescence was observed in the leaves co-expressing MR3_(-)eGFPMut2_, MR3_(-)eGFPMut3_, or MR3_(-)eGFPMut4_ with NP, RdRp_opt_, NSvc4 and VSRs. However, the leaves co-expressing MR3_(-)eGFPMut1_, NP, RdRp_opt_, NSvc4 and VSRs did not show eGFP fluorescence (Fig. 7E). Western blot results agreed with the microscopic observations and showed that eGFP was not expressed from MR3_(-)eGFPMut1_ in cells (Fig. 7F), indicating that deletion of the first 159 aa residues of NS3 significantly affect MR3_(-)eGFP_ mini-replicon to express the eGFP. It is noteworthy that the Northern blot result showed that the expression of gRNA and agRNA from MR3_(-)eGFPMut1_ was not affected (Fig. 7G). We also noticed that the mobility of gRNA and agRNA from MR3_(-)eGFPMut2_ and MR3_(-)eGFPMut4_ was altered compared with that from MR3_(-)eGFPMut1_, MR3_(-)eGFPMut3_ or MR3_(-)eGFP_. Therefore, we conclude that the *NS3* ORF sequence is not necessary for viral replication but is required for the expression of eGFP from the RNA3_(-)_-derived mini-replicons in cells.

RSV RNA2, 3 and 4 IGRs can form secondary hairpin-like structures which is postulated to act as transcription termination signals (24, 25). To further investigate the possible role of RSV NS3 in viral transcription regulation, we predicted the RNA secondary structure and examined the possible RNA-RNA interaction between NS3 coding sequence, IGR and 3′-UTR of RSV RNA3. The 3′-UTR, NS3, and IGR of RSV RNA3 have 65, 636 and 742 nucleotides (nt), respectively. The secondary structure analysis showed that the1-295 nt sequence of IGR formed a very long hairpin structure (Fig. 8). Strikingly, the 1-40 nt coding region sequence of NS3 base-paired with the 575-596 nt sequence of IGR and the 1-65 nt sequence of 3′-UTR and formed a long hairpin structure (Fig. 8). The 619-636 nt coding region sequence of NS3 base-paired with the 396-403 nt and the 537-543 nt sequences of IGR and formed a small hairpin-like structure. The 41-203 nt coding region sequence of NS3 itself formed four long and short hairpin structures. The 205-618 nt coding region sequence of NS3 formed sector structure containing at least 9 hairpins (Fig. 8). The secondary structure analysis suggested that the coding sequence of NS3 likely interacts with IGR and 3′-UTR of RNA3 in forming hairpin-like structure.

**Fig. 8.**
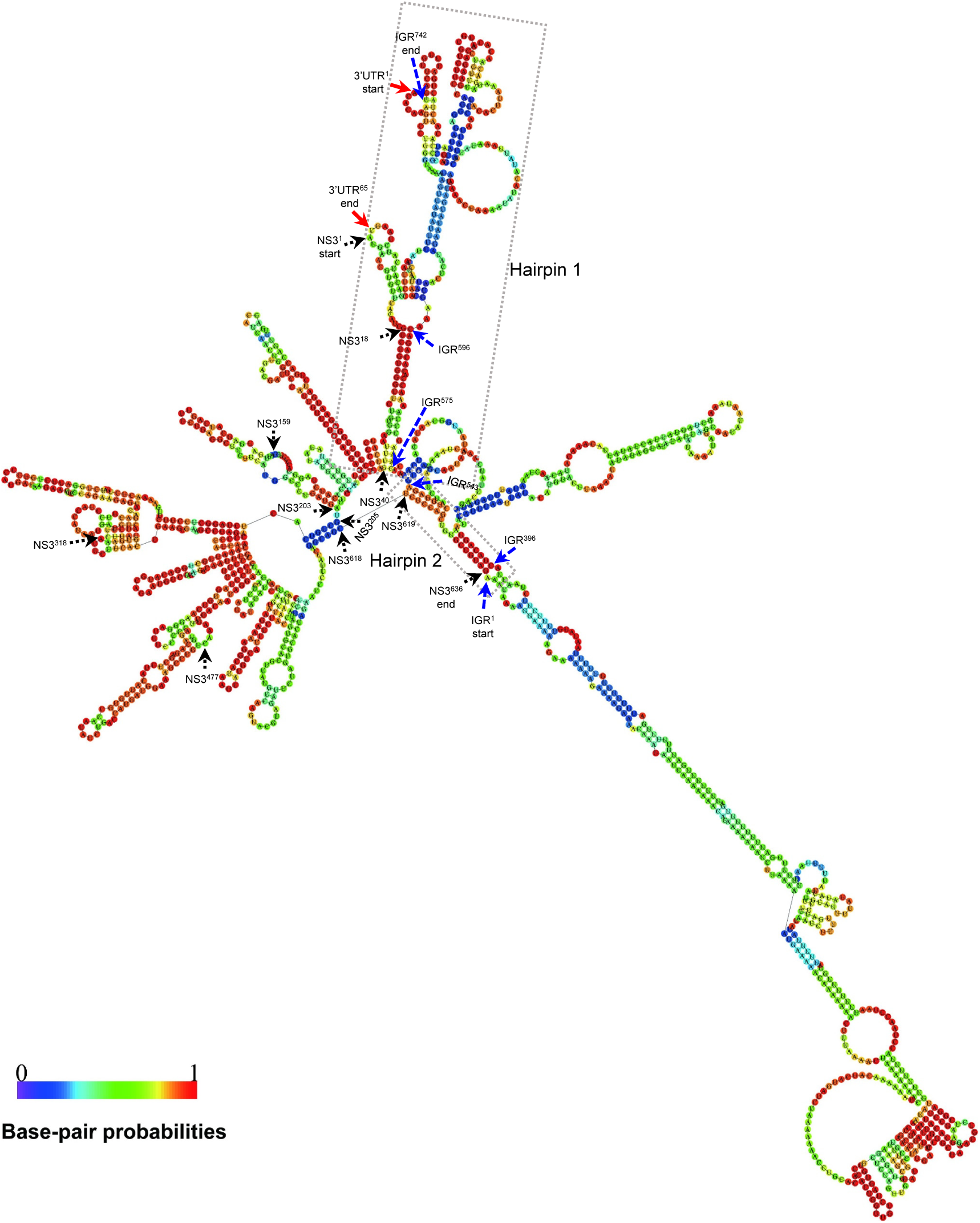
RNA secondary structures of the 3′-UTR, NS3 ORF and IGR region of RSV RNA3 segment. Secondary structure was predicted using the RNA fold web server (http://rna.tbi.univie.ac.at/cgi-bin/RNAWebSuite/RNAfold.cgi) based on thermodynamic prediction of minimal free energy (MFE). Red, black and blue arrows indicated the nucleotide positions in 3′-UTR, NS3 and IGR regions, respectively. Dotted boxes indicated that Hairpin 1 and 2 were formed by IGR, NS3 and 3′-UTR of RSV RNA3, respectively.

### Development of RSV RNA1, RNA2 and RNA4 mini-replicons

RSV genome consists of four RNA segments encoding seven proteins using an antisense or an ambisense coding strategy (15–18). Because the MR3_(-)eGFP_ mini-replicon is functional in *N. benthamiana* leaf cells (Fig. 1 and Fig. 2), we decided to develop mini-replicons for other three RSV genomic RNAs. We first produced MR1_(-)eGFP_ and then replaced the *RdRp* ORF with the *eGFP* gene to produce MR1_(-)eGFP_ (Fig. 9A). For RSV genomic RNA2 and 4, we first cloned the full-length RNA2 or 4 segments individually into the vector, and then replaced the *NSvc2* ORF with the *eGFP* gene to produce MR2_(-)eGFP_ or replaced the *NSvc4* ORF with the *eGFP* gene to produce MR4_(-)eGFP_ (Fig. 9A). *N. benthamiana* leaves were then infiltrated with the mixed Agrobacterium cultures carrying various combinations of plasmids (Fig. 9B). By 5 dpi, strong eGFP fluorescence was observed in the leaves co-expressing MR1_(-)eGFP_, MR2_(-)eGFP_, or MR4_(-)eGFP_ with NP, RdRp_opt_ and four VSRs (Fig. 9B). As expected, the leaves infiltrated with the mixed Agrobacterium cultures lacking NP, RdRp_opt_ or NP+RdRp_opt_ did not show eGFP fluorescence.

**Fig. 9.**
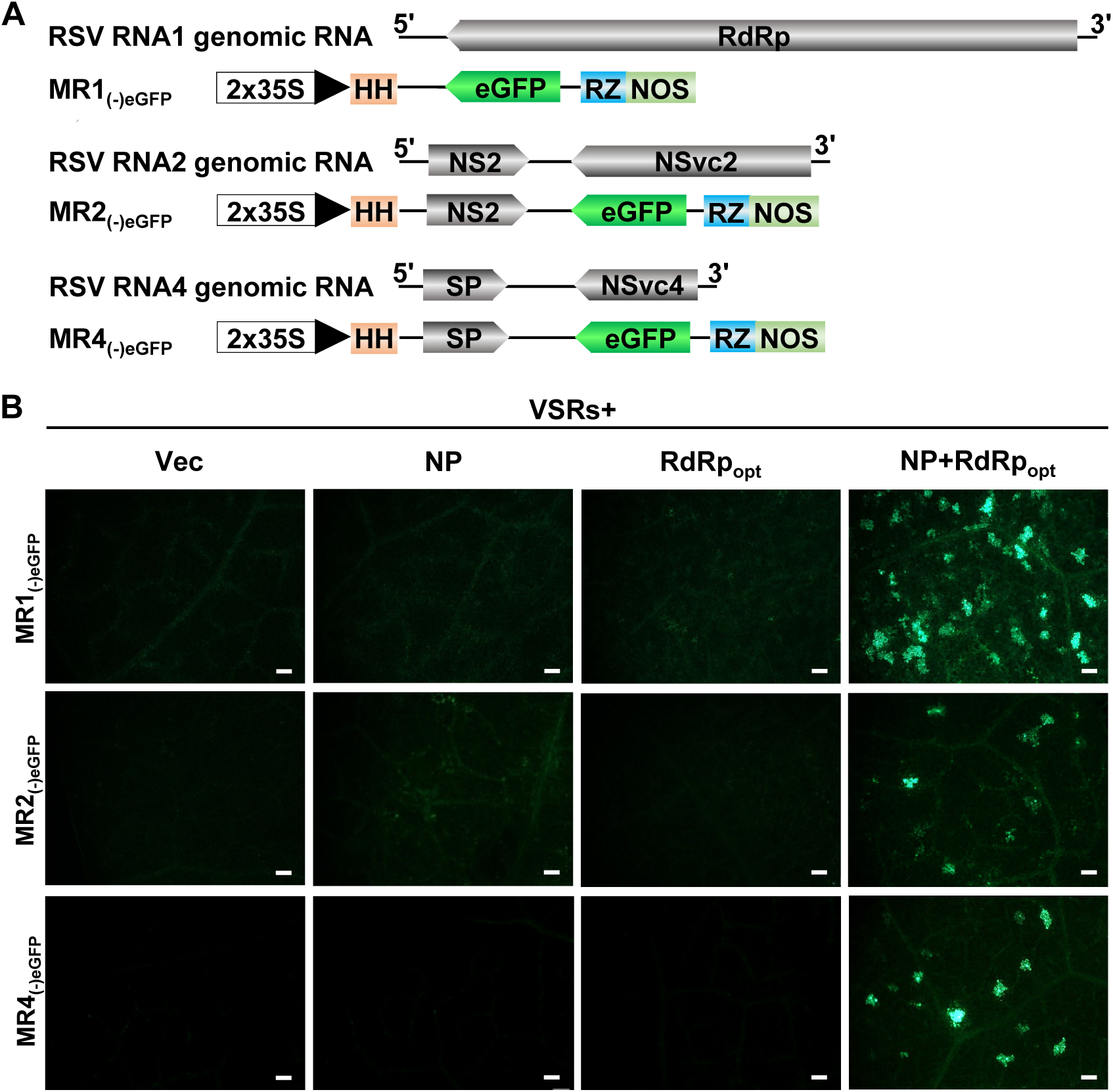
Establishment of complete mini-replicon systems representing RSV RNA1, RNA2, RNA3, and RNA4 genomic RNA segments in *N. benthamiana* leaves. (A) Schematics representing MR1_(-)eGFP_, MR2_(-)eGFP_, and MR4_(-)eGFP_ mini-replicons. The RSV *RdRp* gene in MR1_(-)eGFP_, the *NSvc2* gene in MR2_(-)eGFP_, and the *NSvc4* gene in MR4_(-)eGFP_ were replaced with an *eGFP* gene to produce MR1_(-)eGFP_, MR2_(-)eGFP_, and MR4_(-)eGFP_, respectively. Minus sign (-) and 5′ to 3′ designation represent the viral (genomic)-strand of RNA1, 2 and 4. (B) *N. benthamiana* leaves were infiltrated with various mixed Agrobacterium cultures as described in the figures. The infiltrated leaf tissues were harvested at 5 dpi, and examined and photographed under an inverted fluorescence microscope. Bars = 200 μm.

Collectively, the mini-replicon-based reverse-genetics system, representing all four RSV genomic RNAs, has been created.

## Discussion

There are 3 genera of segmented NSR viruses infecting plants: *Orthotospovirus, Tenuivirus* and *Emaravirus*. The reverse-genetics systems have been established for TSWV and RRV in the genera *Orthotospovirus* and *Emaravirus* (9, 48, 49), respectively. Here, we established a mini-replicon-based reverse-genetics system for RSV, the representative virus for the genus *Tenuivirus.* RSV is an important rice virus and poses significant threat to rice productions in China and many other Asian countries (6, 10, 11). During the past 20 years, the lack of a reliable reverse-genetics system significantly hampered the studies of RSV gene functions and disease induction in plant. To overcome this obstacle, we launched a multiyear research that finally yielded a functional mini-replicon-based reversed genetics system for RSV studies. We first developed a mini-replicon system to express RSV MR3_(-)eGFP_, NP, and a codon usage optimized RdRp (RdRp_opt_), respectively. Using this mini-replicon systems we determined that RSV NP and RdRp_opt_ are indispensable for the eGFP expression from MR3_(-)eGFP_. The expression of eGFP from MR3_(-)eGFP_ was significantly enhanced in the presence of NSs and P19-HcPro-γb. In addition, NSvc4, the movement protein of RSV, facilitated eGFP trafficking between cells. Interestingly, co-expression of RSV NS3 inhibited eGFP expression from MR3_(-)eGFP_. We also found that the RSV *NS3* gene sequence is not necessary for viral replication, but regulates viral RNA expression. The secondary structure analysis showed that the coding sequence of NS3 base-paired with the sequence of IGR and 3′-UTR of RNA3 to form a long hairpin structure. The phenomenon of coding sequence as a *cis*-element in regulating viral RNA expression has not been reported previously for the negative-stranded/ambisense RNA viruses. Finally, based on the system of RNA3_(-)_, we have also produced mini-replicons representing all RSV RNA genomic segments, allowing RSV functional studies in plant.

The choice of promoter for RNA transcriptions is critical for the development of reverse-genetics systems for plant NSR viruses. The genomic and antigenomic RNAs generated from the negative-stranded RNA virus clones were not infectious because their infectious ribonucleoprotein complexes (RNPs) contain not only viral gRNA, but also NP and RdRp proteins (36, 51). Genomic RNAs of the same segmented NSV contain highly conserved 5′ and 3′ terminal untranslated sequences that have eight complementary nucleotides, capable of forming panhandle-like structures. These structures are known to play critical roles in viral gRNA and agRNA replications (51). Moreover, the NSR viruses RNAs do not possess 5′ cap-structures and 3′ poly(A) tails. The classical bacteriophage T7 promoter can produce accurate viral RNA 5′ end sequences without a cap. We initially produced an RSV mini-replicon systems using the T7 promoter. This mini-replicon system, however, did not express the *eGFP* gene from an RNA3-based mini-replicon in plant cells. In our recent report, we also reported that the T7 promoter-based system was unable to generate infectious TSWV RNA transcripts in plant cells (48). It is possible that the synthesis of viral genomic RNA transcripts through the T7 promoter and T7 RNA polymerase is incomplete or is inefficient in *planta*.

In a recent report, we described an expressing vector with a double CaMV 35S promoter (an RNA Pol II promoter), a hammerhead (HH) ribozyme, and an HDV ribozyme to produce infectious TSWV viral RNAs in plant cells (48). The mini-replicon system produced in this study also has an HH ribozyme and an HDV ribozyme before and after the viral sequence to ensure the correct ends. The results shown in Fig. 1 and Fig. 2 suggest that the vector transcribed RSV genomic RNA did bind to viral NP and RdRp to form functional RNPs, which is needed for the synthesis of functional viral gRNA and agRNA. This finding supports earlier reports that the Pol II promoter can not only replicate non-segmented plant NSR viruses (44, 46), but also segmented plant NSR viruses (9, 48).

RSV RdRp is one of the major components needed for the initiation of viral genomic RNA replication (17). RSV RdRp is a 337 kDa protein. When this protein was co-expressed with the mini-replicon in plant cells, no eGFP fluorescence was observed in the infiltrated leaves (Fig. 1A-C). Our computer prediction suggested that the RSV *RdRp* sequence contained many putative cryptic intron splicing sites. Because the segmented plant NSR viruses replicate in cytoplasm, their *RdRp* gene sequences should not been evolved to remove those cryptic intron splicing sites that could be spliced in cell nucleus. We speculated that after the wild-type RSV *RdRp* sequences were expressed, through the 2×35S promoter, in nucleus, they were quickly spliced, resulting in non-functional *RdRp* fragments. After the putative intron splicing sites were removed and the codon usage was optimized, the expressed RdRp_opt_ is capable of supporting *eGFP* expression from the mini-replicon (Fig. 1D). In this study, we also determined that the less concentrated Agrobacterium culture carrying RdRp_opt_ (OD_600_ 0.05) caused higher eGFP expression in cells. In contrast, increase of Agrobacterium culture carrying RdRp_opt_ from OD_600_ 0.1 to OD_600_ 0.4 decreased eGFP expression from the mini-replicon (Fig. 4B and 4C), suggesting that an optimum concentration of RdRp_opt_ is required during RSV infection in plant.

Analyses of the five different VSRs have indicated that in the presence of NSs or P19-HcPro-γb, the eGFP expression from the mini-replicon was significantly enhanced (Fig. 3B and 3C). These VSRs are known to function at different steps in host RNA interference (RNAi) pathway during virus infection in plant (44, 52). Consequently, we conclude that these steps in the RNAi pathway can all affect eGFP expression from the mini-replicon. It is noteworthy that the presence of RSV NS3 alone or NS3 plus one or three of the four VSRs (i.e., NSs or P19-HcPro-γb) suppressed eGFP expression from the RNA3 mini-replicon (3B and 3C) suggesting that RSV NS3 is a negative regulator of RSV RNA3 mini-replicon expression. Similar phenomenon was also reported for TSWV NSs during virus rescue assays using cDNA clones (48). We hypothesize that this negative regulation is caused by the co-suppression of *NS3* gene expression. In this study, addition of NSvc4 significantly increased the number of cells with eGFP fluorescence (Fig. 5A and 5B), further conforming its role in cell-to-cell trafficking.

When the *NS3* gene was replaced with a *mCherry* gene (MR3_(-)mCherry&eGFP_) or with an *eGFP* gene in MR3_(+)eGFP_, no mCherry or eGFP fluorescence was observed in the leaf tissues co-expressing NP, RdRp_opt_ and VSRs (Fig. 6C and 6D). Deletion of the *NS3* gene sequence abolished the function of MR3_(-)eGFP_. However, gRNA and agRNA were detected in the leaves co-expressing RNA3_(+)_, NP, RdRp_opt_, NSvc4 and VSRs (Fig. 6A and 6E). The MR3_(-)eGFP&NS3stop_ mini-replicon contains a translation stop codon immediately after the start codon of NS3 and is still functional in *N. benthamiana* leaves (Fig. 7A, 7B and 7C). This finding suggests that the *NS3* coding sequence is required for the eGFP expression of the mini-replicon, probably required for viral transcription. Through nucleotide deletion assays, we determined that the region encompassing nt 1-477 in the *NS3* ORF is not sufficient to express eGFP from the MR3_(-)eGFP_ mini-replicon (Fig. 7D, and 7E-G), even though the mutant mini-replicon (MR3_(-)eGFPMut1_) is capable of expressing RNA in cells, based on the Northern blot results. Because the positions of the gRNA and agRNA bands from MR_3(-)eGFPMut2_ and MR_3(-)eGFPMut4_ were altered, we speculate that the *NS3* ORF sequence may act as a *cis*-regulatory element during RSV RNA3 viral transcription. Importantly, the secondary structure analysis suggested that the coding sequence of NS3 interacts with IGR and 3′-UTR to form a hairpin-like structure and NS3 itself also forms sophisticated hairpin-like structure (Fig 8). Hairpin structure of TSWV and RSV IGR have been shown to regulate viral transcription termination (53). This is consistent with our findings that *NS3* coding sequence involved in regulation of viral RNA transcription. In our earlier studies, deletion of *NSs* coding sequence from the TSWV S-based mini-replicon system had no clear effect on RNA synthesis (48). Although the genome structure of RSV RNA3 is similar to that of TSWV S RNA, this is the first evidence showing that the *NS3* coding region can act as *cis*-element to regulate the synthesis of viral RNA transcripts of plant segmented NSR viruses.

Based the established mini-replicon system for MR3_(-)eGFP_, we have also produced mini-replicons to express MR1_(-)eGFP_, MR2_(-)eGFP_, and MR4_(-)eGFP_ as described in (Fig. 9A). In the presence of NP, RdRp_opt_ and VSRs, eGFP was expressed from MR1_(-)eGFP_, MR2_(-)eGFP_, and MR4_(-)eGFP_ mini-replicons, respectively (Fig. 9B). We also constructed full-length infectious cDNA clones representing RSV genomic RNA1, RNA2 and RNA4. Infiltration of *N. benthamiana* leaves with the mixed Agrobacterium culture carrying RNA1_(-)_, RNA2_(-)_, RNA3_(-)_, RNA4_(-)_and NP+RdRp_opt_+VSRs did not yield a systemic infection. Because *N. benthamiana* is an experimental host of RSV and the accumulations of RSV RNAs and proteins are lower than that in the rice plants, it is possible that the low level of RSV RNAs and proteins in the infiltrated *N. benthamiana* leaves fails to rescue of RSV systemic infection. It is also possible that the difficulty of delivering four RSV RNA segments into the same cells prevents the systemic infection. We have also infiltrated rice callus tissues with the mixed Agrobacterium culture carrying MR1_(-)eGFP_, MR2_(-)eGFP_, MR3_(-)eGFP_, MR4_(-)eGFP_ and NP+RdRp_opt_+VSRs, none of them work in rice callus tissues.

In conclusion, there are 3 genera of segmented NSR viruses infecting plants and the replicon-based reverse-genetics system has been established for 2 genera. We now established a mini-replicon-based reverse-genetics system for a tenuivirus in *N. benthamiana*. This is the first mini-replicon-based reverse-genetics system for the segmented monocot-infecting tenuivirus, and will provide a useful platform for studies of RSV gene functions during viral replication, cell-to-cell movement and interactions between RSV and host factors in plants. Knowledge learned from this study also benefit the future constructions of full-length multi-segmented infectious clones for tenuivirus in plants.

## Materials and Methods

### Plant growth and virus source

*Nicotiana benthamiana* plants were grown inside a growth chamber maintained at 25°C and a 16/8 h light and dark photoperiod, and used for assays at about 7-leaf stage. RSV was originally isolated from an RSV-infected rice plant as reported (12). Optimization of RSV *RdRp* ORF codon usage and deletion of putative intron splicing sites were performed based on the predictions using the GeneArt^TM^ Project Manager online software (https://www.thermofisher.com/order/geneartgenes/projectmgmt).

### Plasmid construction

#### Constructions of RSV RdRp, RdRp_opt_, NP, NSvc4, NS3 and VSRs mini-replicons

Complementary DNAs (cDNAs) of *RSV*, *NP*, *RdRp*, *RdRp_opt_*, *NS3* and *NSvc4* gene were individually amplified from a total RNA sample isolated from an RSV-infected rice plant through reverse transcription polymerase chain reaction (RT-PCR) using gene specific primers. The resulting RT-PCR products were inserted individually into the expression vector pCambia2300 (refers to as p2300 thereafter), pBINplus, or pCXSN to generate p2300-RdRp_wt_, pBIN-NS3, p2300-NP and pCXSN-NSvc4, respectively. The pCXSV-NSs vector was constructed by inserting a NSs fragment amplified from a cDNA from a TSWV-infected *N. benthamiana* plant into the pCXSN vector. Plasmid pCB301-P19-HcPro-γb (P19-HcPro-γb) that can simultaneously expresses the *Tomato bushy stunt virus* P19 protein, the *Tobacco etch virus* HcPro protein, and the *Barley stripe mosaic virus* γb protein is from a previously published source (46). To construct p2300-RdRp_opt_, we first optimized RdRp_wt_ codon usage and deleted the putative intron splicing sites in it at the GenScript Biotech Corp (Nanjing, China) followed by inserting the synthesized sequence into the p2300 vector to produce p2300-RdR_opt_ (RdRp_opt_).

#### Constructions of MR3_(-)eGFP_, MR3_(-)mCherry&eGFP_ and MR3_(+)eGFP_ mini-replicons

To generate the MR3_(-)eGFP_ and the MR3_(+)eGFP_ mini-replicons, we first prepared cDNAs from a total RNA sample isolated from an RSV-infected rice plant using the Reverse Transcription Kit as instructed (Promega, Madison, WI, USA). We then amplify the full-length RSV RNA3_(-)_ and RNA3_(+)_ sequences from this cDNA through PCR using RSV RNA3 specific primers (Table S1) and a Phanta Super-Fidelity DNA Polymerase (Vazyme Biotech, Nanjing, China). Both PCR products contained a hammerhead (HH) ribozyme (54) before the 5′ end, and then cloned individually behind the 2×35S promoter in the pCB301-2×35S-RZ-NOS vector to generate pCB301-2×35S-HH-RNA3_(-)_-RZ-NOS (refers to as RNA3_(-)_) or pCB301-2×35S-HH-RNA3_(+)_-RZ-NOS (RNA3_(+)_). The presence of an HH and a Hepatitis delta virus ribozyme (RZ) in these two vectors allow the productions of RNA3_(-)_ and RNA3_(+)_ with near perfect ends. To produce a MR3_(-)eGFP_ and a MR3_(+)eGFP_ mini-replicon, we first amplified the *eGFP* gene from SR_(+)eGFP_ (48) using primer FMF37 and FMF38, and then used it to replace the *NP* gene in the RNA3_(-)_ and RNA3_(+)_ through *in vitro* homologous recombination using a ClonExpress II One Step Cloning Kit (Vazyme Biotech, Nanjing, China). The resulting plasmids were referred to as MR3_(-)eGFP_ and MR3_(+)eGFP_, respectively. To produce an MR3_(-)mCherry&eGFP_ min-replicon with both *mCherry* and *eGFP* gene, we PCR-amplified the *mCherry* gene from the TSWV SR_(-)mCherry&eGFP_ mini-replicon as reported previously (48) using primer FMF267 and FMF269, and used it to replace the *NS3* gene in MR3_(-)eGFP_ through homologous recombination as describe above. All primers used in this study are listed in Table S1.

#### Constructions of mutant MR3_(-)eGFP_ mini-replicons

To produce these mutant mini-replicons, we first introduced a stop codon (TAA) after the original start codon of the *NS3* gene through PCR using MR3_(-)eGFP_ as the template, and primer FMF528 and FMF36. This NS3stop fragment was then used to replace the *NS3* gene in MR3_(-)eGFP_ through homologous recombination using prime FMF40 and FMF529. The resulting plasmid is named as MR3_(-)eGFP&NS3stop_. We then deleted a 636 nucleotides (nt) fragment from the *NS3* gene in MR3_(-)eGFP_ through PCR using primer LLY40 and FMF42, and then inserted it into the pCB301 vector through homologous recombination using prime FMF37 and LLY105 to produce MR3_(-)eGFPΔNS3_.

To further investigate the role of RSV *NS3* gene in viral RNA replication and transcription from RNA3_(-)_, we divided *NS3* ORF into four fragments and amplified them individually through PCR using specific primers. These four fragments are: nucleotide position 1-159, 160-318, 319-477, and 478-636, respectively. These fragments were then used to generate MR3_(-)eGFPMut1_, MR3_(-)eGFPMut2_, MR3_(-)eGFPMut3_, and MR3_(-)eGFPMut4_, respectively. Primers used in this study are listed in Table S1. The plasmids were transformed individually into *Agrobacterium tumefaciens* strain GV3101 through electroporation, and the transformants are maintained at -80°C until use.

### Agrobacterium infiltration

*A. tumefaciens* culture were prepared as described previously (44, 45). Briefly, agrobacterium cultures carrying specific plasmids were grown individually in a culture medium and then diluted to OD_600_ = 1.0, or as indicated in the text, in an infiltration buffer (10 mM MES and 10 mM MgCl2, pH 5.6, supplemented with 100 μM acetosyringone). After 2-3 h incubation in the dark and at room temperature (RT), Agrobacterium cultures harboring p2300-NP (OD_600_ = 0.2), p2300-RdRp (OD_600_ = 0.05), p2300-NSs (OD_600_ = 0.1), pCB301-P19-HcPro-γb (OD_600_ = 0.1) or one of the vectors (OD_600_ = 0.2 each) with an *eGFP* and/or an *mCherry* report gene were mixed in equal volumes, or as indicated in the text. The mixed cultures were then infiltrated individually into leaves of 6-7 leaf-stage-old *N. benthamiana* plants using 1 mL needless syringes. The infiltrated plants were grown inside a growth chamber with the same growth conditions as described above.

### Western blot assay

The agro-infiltrated *N. benthamiana* leaves were harvested at 5 days post agro-infiltration (dpi) and homogenized (1 mg/sample) individually in 1 mL extraction buffer (150 mM NaCl, 25 mM Tris-HCl, pH 7.5, 1 mM EDTA, 2% polyvinylpolypyrrolidone, 10 mM dithiothreitol, 10% glycerol, 0.5% Triton X-100, and 1× protease inhibitor cocktail reagent). The crude extract from each sample was mixed with a 5 ×Cloading buffer at a 1:4 ratio (v/v). All the samples were boiled for 10 min and then incubated on ice for 5 min followed by electrophoresis in 12% SDS-PAGE gels. After the protein bands were transferred onto PVDF membranes (GE Healthcare, UK), the blots were probed with an RSV NP specific (1:5000 diluted) or an eGFP specific (1:3000 diluted) antibody followed by a horseradish peroxidase (HRP)-conjugated goat anti-mouse or anti-rabbit secondary antibody (1:10000 diluted). The detection signal was visualized using the ECL Substrate Kit as instructed (Thermo Fisher Scientific, Rockford, USA). The ponceau S-stained Rubisco large subunit gels were used to show sample loadings.

### Northern blot assay

To detect the expressions of RSV gRNAs, agRNAs, and *eGFP* mRNA, we isolated total RNA from the agro-infiltrated *N. benthamiana* leaf tissues using the RNAprep Pure Plant Kit (Tiangen Biotech, Beijing, China). The isolated total RNA samples were separated in 1% formaldehyde agarose gels through electrophoresis, and then transferred onto Hybond-N^+^ membranes (GE Healthcare, UK) (55). DIG-labeled RNA probes specific for the sense or antisense *eGFP* mRNA were *in vitro* synthesized Basel, Switzerland). The blotted membranes were probed with DIG-labeled RNA probes specific for the sense or antisense *eGFP* mRNA. The detection signal was visualized using a DIG-High Prime Detection Starter Kit II as instructed (Roche).

### Fluorescence microscopy

The agro-infiltrated *N. benthamiana* leaf tissues were harvested at 5 dpi, and examined under an inverted fluorescence microscope (OLYMPUS IX71-F22FL/DIC, Tokyo, Japan) equipped with a green barrier filter. The captured images were processed using the ImagePro system (OLYMPUS, Tokyo, Japan) and then the Adobe Photoshop CS4 (San Jose, CA, USA).

## Acknowledgments

This work was supported by the National Natural Science Foundation of China (31630062, 31925032 and 31870143), the Fundamental Research Funds for the Central Universities (JCQY201904 and KYXK202012), Youth Science and Technology Innovation Program to XT, Project funded by China Postdoctoral Science Foundation (2020M681643) to MF.

## Author contributions

M.F., L.L. and X.T. conceived and designed the experiments and M.Y., Y.Z., J.W. and Y.X. provided input. M.F., L.L., R.C., Y.Y., Y.D., M.C., and G.R. performed the experiments. M.F., X.D., X.Z. and X.T. wrote the manuscript.

## Data availability

All data produced in this study are presented in this manuscript or as the supporting files.

---

Table S1. List of primers used in the study.

Table S2. The predicted intron splicing sites of wild-type RdRp gene.

Table S3 Optimized RdRp gene sequence made in this study.

