## Supplemental Table 1, 2 and 3 for "Development of a mini-replicon-based reverse-genetics system for rice stripe tenuivirus"

### Table S1. List of primers used in the study.

| Construct | Abbreviation | Primer sequence (5' to 3') |
|---|---|---|
| p2300-NP | NP | **F:** GAGCTCATGGGTACCAACAAGCCAG<br>**R:** GGATCCCTAGTCATCTGCACCTTCTGC |
| p2300-RdRp$^{wt}$ | RdRp$^{wt}$ | **F:** GAGAGGTCGCGAGCTCGGTACCATGACGACACCACCTCTCGTTATACC<br>**R:** GCATGCCTGCAGGTCGACTCTAGATCAGAAATCGAACTTATGGTC |
| p2300-RdRp$^{opt}$ | RdRp$^{opt}$ | **F:** GAGAGGTCGCGAGCTCGGTACCATGACTACCCCTCCTCTTGTG<br>**R:** GCATGCCTGCAGGTCGACTCTAGACTAGAAGTCGAACTTGTGATC |
| pCXSN-NSs | NSs | **F:** CTCGGTACCATGTCTTCAAGTGTTTATGAG<br>**R:** GACTCTAGATTATTTTGATCCTGAAGCATATG |
| pBIN-NS3 | NS3 | **F:** CTCGGTACCATGAACGTGTTCACATCGTC<br>**R:** AGTCTAGACTACAGCACAGCTGGAGAGC |
| pCXSN-NSvc4 | NSvc4 | **F:** CTCGGTACCATGGCTTTGTCTCGACTTTTG<br>**R:** GACTCTAGACTACATGATGACAGAAACTTC |
| pCB301-HH-RNA1$_{(-)}$-RZ-NOS | RNA1$_{(-)}$ | **F:** GGAATATCTTCCTATAGTCACACAAAGTCCAGAGGAAAACAAAATG<br>**R:** GTGGAGATGCCATGCCGACCCACACATAGTCAGAGGAAAAAATAATTTTG<br>**F**[a]: CAAAATTATTTTTTCCTCTGACTATGTGTGGGTCGGCATGGCATCTCCAC<br>**R**[a]: CATTTTGTTTTCCTCTGGACTTTGTG TGACTATAGGAAGATATTCC |
| pCB301-HH-RNA2$_{(-)}$-RZ-NOS | RNA2$_{(-)}$ | **F:** GGAATATCTTCCTATAGTCACACAAAGTCCTGGGTATATAAGCCAC<br>**R:** GTGGAGATGCCATGCCGACCCACACAAAGTCTGGGTATAACTTCTTCG<br>**F**[a]: GAAGAAGTTATACCCAGACTTTGTGTGGGTCGGCATGGCATCTCCAC<br>**R**[a]: GTGGCTTATATACCCAGGACTTTGTGTGACTATAGGAAGATATTCC |
| pCB301-HH-RNA3$_{(-)}$-RZ-NOS | RNA3$_{(-)}$ | **F:** CGAAACTATAGGAATATCTTCCTATAGTCACACAAAGTCCTGGGTAAAATAGTTAT<br>**R:** GGTGGAGATGCCATGCCGACCCACACAAAGTCTGGGTAATAAAATTTTC<br>**F**[a]: GAAAATTTTATTACCCAGACTTTGTGTGGGTCGGCATGGCATCTCCACC<br>**R**[a]: GTTTCGGCCTTTCGGCCTCATCAGACACAAAGAGGCCTCTCCAAATGAAATG |
| pCB301-HH-RNA3$_{(+)}$-RZ-NOS | RNA3$_{(+)}$ | **F:** CGAAACTATAGGAATATCTTCCTATAGTCACACAAAGTCTGGGTAATAAAATTTTC<br>**R:** GAGGTGGAGATGCCATGCCGACCCACACAAAGTCCTGGGTAAAATAGTTATATTTTTAC<br>**F**[a]: CTATTTTACCCAGGACTTTGTGTGGGTCGGCATGGCATCTCCACC<br>**R**[a]: GAAAATTTTATTACCCAGACTTTGTGTGACTATAGGAAGATATTCCTATAGTTTCG |
| pCB301-HH-RNA4$_{(-)}$-RZ-NOS | RNA4$_{(-)}$ | **F:** GGAATATCTTCCTATAGTCACACAAAGTCCAGGGCATTTGTACAACG<br>**R:** GGTGGAGATGCCATGCCGACCCACACAAAGTCAGGGCATATCTTTTGAG<br>**F**[a]: CTCAAAAGATATGCCCTGACTTTGTGTGGGTCGGCATGGCATCTCCACC<br>**R**[a]: CGTTGTACAAATGCCCTGGACTTTGTGTGACTATAGGAAGATATTCC |
| pCB301-HH-RNA3$_{(-)eGFP}$-RZ-NOS | MR3$_{(-)eGFP}$ | **F:** CCACAACTACTGATTTGTTCAGTTTACTTGTACAGCTCGTCCATGCCGAGA<br>**R:** CATTCCTCCAGTACCTCTTGCTAGAATGGTGAGCAAGGGCGAGGAGCTGTTC<br>**F**[a]: GAACAGCTCCTCGCCCTTGCTCACCATTCTAGCAAGAGGTACTGGAGGAATG<br>**R**[a]: TCTCGGCATGGACGAGCTGTACAAGTAAACTGAACAAATCAGTAGTTGTGG |
| pCB301-HH-RNA3$_{(+)eGFP}$-RZ-NOS | MR3$_{(+)eGFP}$ | **F:** CATTCCTCCAGTACCTCTTGCTAGAATGGTGAGCAAGGGCGAGGAGCTGTTC<br>**R:** CCACAACTACTGATTTGTTCAGTTTACTTGTACAGCTCGTCCATGCCGAGA<br>**F**[a]: TCTCGGCATGGACGAGCTGTACAAGTAAACTGAACAAATCAGTAGTTGTGG<br>**R**[a]: GAACAGCTCCTCGCCCTTGCTCACCATTCTAGCAAGAGGTACTGGAGGAATG |
| pCB301-HH-RNA3$_{(-)mCherry\&eGFP}$-RZ-NOS | MR3$_{(-)mCherry\&eGFP}$ | **F:** CAATACAATTCCGACATCATCTAAGTATGGTGAGCAAGGGCGAGGAGGATAAC<br>**R:** TCTTTTCTTTTTTTCTTTTTCCTTTTATTTTTAAGATCTGTACAGCTCGTCCATGC<br>**F**[a]:GCATGGACGAGCTGTACAGATCTTAAAAATAAAAGGAAAAAGAAAAAAAGAAAAG<br>**R**[a]: GTTATCCTCCTCGCCCTTGCTCACCATACTTAGATGATGTCGGAATTGTATTG |

| | | |
|---|---|---|
| pCB301-HH-RNA3$_{(-)eGFP\Delta NS3}$-RZ-NOS | MR3$_{(-)eGFP\Delta NS3}$ | **F:** AAATAAAAGGAAAAAGAAAAAAAGAAAAGAAAACAAATAATC |
| | | **R:** GGTGGAGATGCCATGCCGACCCACACAAAGTCTGGGTAATAAAATTTTC |
| | | **F[a]:** GAAAATTTTATTACCCAGACTTTGTGTGGGTCGGCATGGCATCTCCACC |
| | | **R[a]:** TTTTCTTTTTCCTTTTATTTACTTAGATGATGTCGGAATTGTATTG |
| pCB301-HH-RNA3$_{(-)eGFP\&NS3stop}$-RZ-NOS | MR3$_{(-)eGFP\&NS3stop}$ | **F:** CAATTCCGACATCATCTAAGTATGTAAGTGTTCACATCGTCTG |
| | | **R:** GGTGGAGATGCCATGCCGACCCACACAAAGTCTGGGTAATAAAATTTTC |
| | | **F[a]:** GAAAATTTTATTACCCAGACTTTGTGTGGGTCGGCATGGCATCTCCACC |
| | | **R[a]:** GACGATGTGAACACTTACATACTTAGATGATGTCGGAATTG |
| pCB301-HH-RNA3$_{(-)eGFPMut1}$-RZ-NOS | MR3$_{(-)eGFPMut1}$ | **F:** CAATACAATTCCGACATCATCTAAGTAAATTTGACAATAGG |
| | | **R:** TCTTTTCTTTTTTTCTTTTTCCTTTTATTTCTACAGCACAGCTGGAGAGC |
| | | **F[a]:** GCTCTCCAGCTGTGCTGTAGAAATAAA AGGAAAAAGAAAAAAAGAAAAGA |
| | | **R[a]:** ACTTAGATGATGTCGGAATTGTATTGTATAGTAAAAATA |
| pCB301-HH-RNA3$_{(-)eGFPMut2}$-RZ-NOS | MR3$_{(-)eGFPMut2}$ | **F:** CAATACAATTCCGACATCATCTAAGTTTCTTCACTGAAGTGAAGCC |
| | | **R:** TCTTTTCTTTTTTTCTTTTTCCTTTTATTTGACAGGCTTTCCATCATGGTCAG |
| | | **F[a]:** AAATAAAAGGAAAAAGAAAAAAAGAAAAGAAAACAAATAATC |
| | | **R[a]:** GGCTTCACTTCAGTGAAGAAACTTAGATGATGTCGGAATTGTATTG |
| pCB301-HH-RNA3$_{(-)eGFPMut3}$-RZ-NOS | MR3$_{(-)eGFPMut3}$ | **F:** CAATACAATTCCGACATCATCTAAGTATTGATGAGCATCAG |
| | | **R:** TCTTTTCTTTTTTTCTTTTTCCTTTTATTTTTTCACATAAGAGGATGACATC |
| | | **F[a]:** AAATAAAAGGAAAAAGAAAAAAAGAAAAGAAAACAAATAATC |
| | | **R[a]:** ACTTAGATGATGTCGGAATTGTATTGTATAGTAAAAATA |
| pCB301-HH-RNA3$_{(-)eGFPMut4}$-RZ-NOS | MR3$_{(-)eGFPMut4}$ | **F:** CAATACAATTCCGACATCATCTAAGTATGAACGTGTTCAC |
| | | **R:** TCTTTTCTTTTTTTCTTTTTCCTTTTATTTGGAGGGGTGCC |
| | | **F[a]:** AAATAAAAGGAAAAAGAAAAAAAGAAAAGAAAACAAATAATC |
| | | **R[a]:** ACTTAGATGATGTCGGAATTGTATTGTATAGTAAAAATA |
| pCB301-HH-RNA1$_{(-)eGFP}$-RZ-NOS | MR1$_{(-)eGFP}$ | **F:** GTCCCTTTGTTGAAGAGGACTTCTTACTTGTACAGCTCGTCCATGCCGAGA |
| | | **R:**GATTTTGTTTTCCACAAAAGAATTGAAGGATGGTGAGCAAGGGCGAGGAGCTGTTC |
| | | **F[a]:** GAACAGCTCCTCGCCCTTGCTCACCATCCTTCAATTCTTTTGTGGAAAACAAAATC |
| | | **R[a]:** TCTCGGCATGGACGAGCTGTACAAGTAAGAAGTCCTCTTCAACAAAGGGAC |
| pCB301-HH-RNA2$_{(-)eGFP}$-RZ-NOS | MR2$_{(-)eGFP}$ | **F:** CATACATGAATGAACCTATTGGCTTACTTGTACAGCTCGTCCATGCCGAGA |
| | | **R:** GTCTGGGTATAACTTCTTCGAAGATGGTGAGCAAGGGCGAGGAGCTGTTC |
| | | **F[a]:** GAACAGCTCCTCGCCCTTGCTCACCATCTTCGAAGAAGTTATACCCAGAC |
| | | **R[a]:** TCTCGGCATGGACGAGCTGTACAAGTAAGCCAATAGGTTCATTCATGTATG |
| pCB301-HH-RNA4$_{(-)eGFP}$-RZ-NOS | MR4$_{(-)eGFP}$ | **F:** CATACTCCGGAACTGGTTATCTCACTTACTTGTACAGCTCGTCCATGCCGAGA |
| | | **R:** GATTAAGCTAATATATACTTTAATTATGGTGAGCAAGGGCGAGGAGCTGTTC |
| | | **F[a]:** GAACAGCTCCTCGCCCTTGCTCACCATAATTAAAGTATATATTAGCTTAATC |
| | | **R[a]:** TCTCGGCATGGACGAGCTGTACAAGTAAGTGAGATAACCAGTTCCGGAGTATG |
| pGEM-eGFP | - | **F:** ATGGTGAGCAAGGGCGAGGAGCTGTTC |
| | | **R:** TTACTTGTACAGCTCGTCCATGCCGAGA |
| pGEM-anti-eGFP | - | **F:** TTACTTGTACAGCTCGTCCATGCCGAGA |
| | | **R:** ATGGTGAGCAAGGGCGAGGAGCTGTTC |
| pGEM-NS3 | - | **F:** CATGGCGGCCGCGGGAATTCGATTATGAACGTGTTCACATCGTC |
| | | **R:** CAGGCGGCCGCGAATTCACTAGTGATCTACAGCACAGCTGGAGAGC |
| pGEM-antiNS3 | - | **F:** CATGGCGGCCGCGGGAATTCGATTCTACAGCACAGCTGGAGAGC |
| | | **R:** CAGGCGGCCGCGAATTCACTAGTGATATGAACGTGTTCACATCGTC |

[a] Forward and reverse primers were used to amplified the linearized pCB301 vectors by PCR.

**Table S2. The predicted intron splicing sites of wild-type RdRp gene.**

| Position (bp) | | | | | | |
|---|---|---|---|---|---|---|
| 40 | 1233 | 2741 | 4107 | 5500 | 6655 | 7576 |
| 114 | 1258 | 2747 | 4113 | 5550 | 6678 | 7614 |
| 121 | 1275 | 2806 | 4195 | 5607 | 6682 | 7633 |
| 235 | 1293 | 2865 | 4246 | 5737 | 6708 | 7664 |
| 268 | 1313 | 2983 | 4378 | 5771 | 6750 | 7683 |
| 271 | 1369 | 3088 | 4558 | 5784 | 6817 | 7684 |
| 298 | 1385 | 3171 | 4567 | 5887 | 6829 | 7698 |
| 361 | 1536 | 3183 | 4612 | 5931 | 6930 | 7699 |
| 477 | 1558 | 3204 | 4624 | 6055 | 6944 | 7774 |
| 647 | 1654 | 3281 | 4662 | 6076 | 6948 | 7825 |
| 686 | 1675 | 3289 | 4692 | 6099 | 6961 | 7898 |
| 771 | 1767 | 3319 | 4738 | 6139 | 6972 | 7953 |
| 829 | 1806 | 3390 | 4783 | 6160 | 6998 | 7959 |
| 855 | 1861 | 3458 | 4801 | 6163 | 7034 | 8068 |
| 857 | 1887 | 3589 | 4869 | 6180 | 7179 | 8096 |
| 1015 | 1944 | 3640 | 4970 | 6228 | 7255 | 8113 |
| 1018 | 1965 | 3706 | 5023 | 6255 | 7307 | 8134 |
| 1033 | 1977 | 3712 | 5124 | 6360 | 7352 | 8187 |
| 1068 | 2114 | 3784 | 5180 | 6364 | 7457 | 8317 |
| 1087 | 2233 | 3791 | 5215 | 6399 | 7458 | 8346 |
| 1131 | 2353 | 3828 | 5250 | 6412 | 7482 | 8513 |
| 1132 | 2365 | 3940 | 5251 | 6426 | 7486 | 8573 |
| 1163 | 2385 | 3984 | 5277 | 6469 | 7509 | 8595 |
| 1164 | 2398 | 4014 | 5292 | 6476 | 7517 | 8648 |
| 1171 | 2437 | 4066 | 5386 | 6510 | 7546 | 8682 |
| 1176 | 2568 | 4090 | 5440 | 6555 | 7552 | 8704 |
| 1180 | 2709 | 4106 | 5498 | 6576 | 7568 | |

The putative 188 intron splicing sites (ISSs) of wild-type RdRp gene sequence were predicted by Alternative Splice Site Predictor (ASSP) (http://wangcomputing.com/assp/).

**Table S3. Optimized RdRp gene sequence used in the study.**

ATGACTACCCCTCCTCTTGTGATTCCTCTTCATGTGCACGGTAGGTCTTACGAGCTTCTTGCTGGTTACCATGAGGTGGACTG

GCAAGAGATTGAGGAACTGGAAGAGACTGATGTGAGAGGTGATGGCTTCTGCCTGTACCACTCTATCCTTTACTCTATGGGC

CTGAGCAAAGAGAACTCTAGGACCACCGAGTTCATGATCAAGCTGAGGTCTAACCCTGCTATCTGCCAGCTGGATCAAGAG

ATGCAGCTGTCTCTTATGAAGCAGCTGGACCCGAACGATTCTTCTGCTTGGGGTGAAGATATCGCCATCGGCTTCATGGCTAT

CATCCTGCGGATCAAGATTATCGCCTACCAGACCGTGGATGGCAAGCTGTTCAAGACTATCTACGGTGCCGAGTTCGAGAGC

ACCATCAGGATTAGGAACTACGGCAACTACCACTTCAAGAGCCTCGAGACTGATTTCGACCACAAGGTGAAGCTCCGGTCC

AAGATTGAAGAGTTCCTTCGGATGCCTGTTGAGGACTGCGAGTCTATTTCTCTGTGGCACGCTTCTGTGTACAAGCCGATCG

TGTCTGATTCTCTGAGCGGCCACAAGAGCTTCAGCAATGTGGATGAGCTGATCGGCAGCATCATCAGCAGCATGTACAAGAT

CATGGACAACGGCGATCAGTGCTTCCTGTGGTCTGCTATGAGAATGATCGCTCGGCCTTCTGAGAAGCTTTACGCTCTTGCT

GTGTTCCTGGGCTTCAACCTGAAGTTCTACCATGTGAGAAAGCGGGCAGAGAAGCTTACCGCTAAGTTGGAGTCTGATCAC

ACCAACCTGGGTGTGAAGCTTATCGAGGTTTACGAGGTGTCAGAGCCTACTAGGTCTACTTGGGTTTTGAAGCCTGGTGGCT

CTCGGATTACTGAGACTAGGAACTTCGTGATCGAAGAGATCATCGATAACCGGCGGAGCCTTGAGTCTCTGTTCGTGTCATC

TTCTAACTACCCTGCCGAGCTGTGCTCTCAGAAGCTTTCTGCTATCAAGGACCGGATCGCTCTGATGTTCGGCTTTATTAACA

GGACCCCTGAGAACAGCGGTCGTGAGCTTTACATCAACACCTACTACCTGAAGCGGATCCTCCAGGTGGACAGAAATGTGA

TCAGGGACTCTCTTAGGTCACAGCCTGCTGTGGGTATGATCCAGATTATTAGGCTGCCTACCGCTTTCGGCACTTACAATCCT

GAGGTTGGGACCCTTTTGCTTGCTCAGACTGGTCTTATCTACAGGCTTGGTACTACTACCCGGGTGCAGATGGAAGTTAGAA

GGTCCCCATCTGTGATCAGCCGGTCACACAAGATCACTAGCTTCCCCGAGACTCAGAAGCACAACAACAACCTGTACGACT

ACGCTCCGAGGACTCAAGAGACTTTCTACCATCCTAACGCCGAGATCTACGAGGCTGTGGATGTTAAGACCCCTTCCGTGAT

TACCGAGATCGTCGACAACCACATCGTGATTAAGCTGAACACCGACGACAAAGGCTGGTCCGTGAGCGATTCTATCAAGCA

GGATTTCGTGTACCGGAAGCGGCTGATGGACGCTAAGAATATTGTGCACGACTTCGTGTTCGACATCCTGTCTACCGAGACT

GACAAGTCCTTCAAGGGTGCTGATCTGAGCATCGGCGGTATCTCTGATAACTGGTCCCCTGACGTGATCATCAGCCGTGAAT

CTGATCCACAGTACGAGGACATTGTGGTGTACGAGTTCACTACCAGGTCCACCGAGTCTATCGAGTCTCTTCTGAGATCCGT

CGAGGTGAAGTCCCTGAGATACAAAGAGGCTATCCAAGAGAGGGCTATCACCCTGAAGAAGAGGATCAGCTACTACACCAT

CTGCGTGAGCCTTGATGCTGTGGCTACCAACCTTCTTTCTCTGCCTGCTGATGTGTGCAGAGAGCTTATCATTAGGCTCAGGG

TTGCCAACCAGGTGAAGATTCAGCTGGCTGACAACGACATCAACCTGGATTCTGCTACCCTTCTGGCTCCTGACATCTACCG

TATCAAAGAGATGTTCCGAGAGAGCTTCCCGAACAACAAGTTCATTCACCCGATCACCAAAGAAATGTACGAGCACTTCGT

GAACCCCATGATCAGCGGCGAGAAGGATTATGTGGCTAACCTCAAGTCCATCATCGACAAAGAGACTAGGGACGAGCAGCG

GAAGAACCTCGAGTCTTTGAAGGTTGTGGACGGGAAGAAGTACACCGAGAGGAAGGCTGAAACTGCCCTGAATGAGATGT

CTCAGGCTGAGGAACACTACCGGTCTTACTTCGAGAACGACAACTTCCGGTCTACCTTGAAGGCTCCTGTTCAGCTGCCTCT

GATCATCCCTGATGTGTCCTCTCAGGACAACCAGTTCAGCAACAAAGAGCTGAGCGACCGGATCCGTAAGAAGCCTATTGAT

CACCCCATCTACAACATCTGGGACCAAGCTGTGAACAAGCGGAACTGCTCTATTGCTCTTGGTCACCTGGATGAGTTGGAGA

TCTCTATGCTTGAGGGCCAAGTGGCCAAGAAGGTGGAAGAGTCTTACAAGAAGGACCGGTCACAGTACAACAGGACCACC

TTGCTTACCAACATGAAGGAAGATATCTACCTGGCCGAGAGGGGCATCAATGCTAAGAAGAGGCTTGAGGAACCGGACGTC

AAGTTCTACAGGGATCAAAGCAAGCGGCCTTTCCATCCTTTCGTGTCAGAGACAAGGGACATCGAGCAGTTCACTCAGAAA

GAATGCCTCGAGCTGAACGAAGAGTCTGGTCACTGCTCTCTCATCAACGTCGAGGATCTTGTGCTGTCTGCTCTTGAGCTTC

ACGAGGTTGGAGATTTGGAGCACCTGTGGAACAACATCAAGGCCCACTCTAAGACCAAGTTCGCTCTGTACGCCAAGTTCA

TCTCTGATCTTGCTACCGAGCTGGCCATCTCTCTGTCTCAGAATTGCAAAGAGGACACCTACGTCGTCAAGAAGCTGAGGGA

CTTCTCTTGCTACGTGCTGATCAAGCCTGTGAACCTGAAGAGTAACGTGTTCTTCTCCCTGTACATCCCGTCCAACATCTACA

AGTCTCACAACACTACCTTCAAGACCCTGATCGGCTCTCCTGAGTCTGGTTACATGACCGATTTCGTGAGCGCCAACGTGAG

CAAGCTTGTGAATTGGGTTAGATGCGAGGCTATGATGCTGGCCCAAAGAGGATTTTGGAGAGAGTTCTACGCTGTCGCCCCT

TCTATCGAAGAACAGGATGGTATGGCTGAGCCAGAGTCTGTTTGCCAGATGATGTCTTGGACCCTTCTGATCCTGCTGAACG

ATAAGCACCAGCTTGAAGAGATGATCACCGTGTCCAGATTCGTCCACATGGAAGGCTTCGTTACTTTCCCTGCTTGGCCTAA

GCCGTACAAGATGTTCGACAAGCTGTCTGTGACTCCGAGGTCTAGACTTGAGTGCCTTGTGATCAAGCGGCTCATCATGCTG

ATGAAGCACTACAGCGAGAACCCGATCAAGTTTATGATCGAGGACGAGAAGAAGAAGTGGTTCGGCTTCAAGAATATGTTC
CTGCTGGACTGCAACGGCAAGCTCGCTGATCTTTCTGATCAGGACCAGATGCTGAACCTGTTCTACCTGGGTTACCTGAAGA
ACAAGGACGAAGAGGTCGAGGATAACGGTATGGGTCAGCTTCTGACCAAGATCCTGGGATTCGAGTCAGCTATGCCTAAGA
CCAGGGATTTCCTGGGCATGAAGGATCCTGAGTACGGCACCATTAAGAAGCACGAGTTCTCCATCAGCTACGTGAAGGATCT
GTGCGACAAGTTCCTGGACAGGCTGAAGAAAACCCACGGCATTAAGGACCCTATTACCTACCTGGGCGACAAGATCGCTAA
GTTCTTGAGCACCCAGTTCATCGAGACTATGGCTAGCCTGAAGGCCAGCTCTAACTTCAGCGAGGATTACTACCTGTACACC
CCTTCTAGGCGGCTCAAGAATCAAGAACAGAGCCGGTCTAAGCACGTGATCGATGCTGGTGGTAACATCAGCGCTTCAGTG
AAGGGTAAGCTGTACCACAGGTCTAAGGTGATCGAGAAGCTCACTACCCTGATCAAGGATGAGACTCCGGGCAAAGAACTG
AAGATCGTGGTGGATCTTCTGCCGAAGGCTATGGAAGTGCTGAACAAGAACGAGTGCATGCATATCTGCATCTTCAAGAAGA
ACCAGCACGGCGGTCTGCGAGAGATCTATGTGCTTAACATCTTCGAGCGGATCATGCAAAAGACCGTCGAGGACTTCAGCA
GGGCTATTCTTGAGTGTTGCCCGTCTGAGACTATGACCTCTCCAAAGAACAAGTTTCGGATCCCCGAGCTGCACAACATGGA
AGCTAGAAAGACCCTCAAGAATGAGTATATGACCATCAGCACCAGCGACGACGCCTCTAAGTGGAATCAGGGACATTACGT
GTCAAAGTTTATGTGCATGCTCCTGAGGCTGACCCCGACTTACTACCATGGTTTTTTGGTTCAGGCTCTGCAGCTGTGGCACC
ACAAGAAGATTTTCTTGGGAGATCAGCTGCTGCAGCTTTTCAACCAGAACGCCATGCTTAACACCATGGACACCACCTTGAT
GAAGGTGTTCCAGGCTTACAAGGGCGAGATTCAGGTTCCATGGATGAAGGCTGGCCGTAGCTATATCGAAACCGAGACAGG
TATGATGCAGGGCATTCTGCACTACACCTCTTCTCTGTTCCACGCCATTTTTCTGGACCAGCTTGCTGAAGAGTGCCGGCGTG
ATATCAACCGGGCTATTAAGACCATCAACAACAAAGAAAACGAGAAGGTGAGCTGCATCGTGAACAATATGGAAAGCTCCG
ACGACAGCAGCTTCATCATCTCTATCCCGAACTTCAAAGAGAATGAGGCTGCCCAGCTTTACCTTCTGTGCGTGGTGAATAG
CTGGTTTCGGAAGAAAGAGAAGCTCGGCACCTACCTCGGTATCTACAAGAGCCCTAAGTCTACCACTCAGACCCTGTTCGT
GATGGAATTCAACAGCGAGTTTTTCTTCTCCGGCGACGTGCACAGGCCTACTTTTAGATGGGTTAACGCTGCTGTGCTCATCG
GCGAGCAAGAAACCCTTTCTGGCATTCAAGAGGAACTGAGCAACACCCTCAAGGACGTTATCGAAGGTGGTGGAACCTAC
GCTCTTACCTTCATTGTGCAGGTTGCCCAGGCAATGATCCACTACAGAATGCTCGGTTCCAGCGCTTCTTCTGTTTGGCCTGC
TTACGAGACTCTGCTGAAGAACTCTTACGATCCTGCTCTGGGCTTCTTCCTGATGGATAATCCTAAGTGCGCTGGTCTGCTGG
GGTTCAATTACAACGTTTGGATTGCCTGCACCACCACTCCTCTTGGTGAGAAGTACCACGAGATGATCCAAGAGGAAATGAA
GGCAGAGTCCCAGTCTCTGAAGTCTGTTACCGAGGACACCATTAACACCGGTCTTGTGTCTCGGACCACCATGGTTGGTTTC
GGCAACAAGAAACGGTGGATGAAGCTCATGACCACTCTCAACCTTAGCGCTGACGTGTACGAGAAGATCGAGGAAGAACC
GCGGGTCTACTTTTTCCATGCTGCTACTGCTGAGCAGATCATCCAGAAAATCGCTATCAAGATGAAGTCACCGGGCGTGATCC
AGAGCCTTTCTAAGGGTAATATGCTCGCCCGGAAGATCGCCTCTTCAGTGTTCTTCATTAGCCGGCACATCGTGTTCACCATG
AGCGCTTATTACGATGCTGACCCTGAGACACGGAAAACCAGCCTGCTTAAAGAGCTAATCAACAGCTCTAAGATCCCGCAG
AGGCACGATTACTTGCAAGAGCCACATACTCTCAAGCCGACCAAGGTTGAAGTTGACGAGGATAGCTGGGAGTTCAAGAGC
GCAAAAGAGGAATGCATCCGGGTTCTCAAGCAGCGGATTAAGATTCACACTGGCCGGGAAGAGAGATCCATCTCCTTGCTT
TTTGAGAACATGGCCAAGAGCATGATCGGTAGGTGCACCGATCAGTACGATGTGAGGGAAAACGTGTCCATTCTTGCTTGCG
CCCTGAAGATGAACTACTCCATCTTTAAGAAGGACGCTGCCCCGAACAGATACCTGTTGGATGAGAAGAACCTGGTGTACCC
GCTGATCGGTAAAGAGGTTAGCGTCTACGTCAAGAGCGACAAGGTGCACATTGAGATCTCCGAGAAGAAAGAACGGCTGA
GCACCAAGTTGTTCAACATCGATAAGATGAAGGATATCGAGGAAACCCTGTCTCTGCTGTTCCCAAGCTACGGTGATTACCT
GAGCCTGAAAGAGACAATCGACCAGGTCACATTCCAGAGCGCTATCCACAAGGTTAACGAGAGAAGGCGTGTTAGGGCTG
ATGTGCATCTTACTGGTACAGAGGGCTTCTCCAAGCTGCCTATGTATACTGCTGCTGTGTGGGCCTGGTTCGACGTTAAGACT
ATTCCTGCTCACGACAGCATCTACCGGACCATTTGGAAGGTGTACAAAGAGCAGTACTCCTGGCTGAGCGATACCCTGAAAG
AAACCGTTGAGAAGGGGCCTTTCAAGACCGTTCAGGGCGTTGTGAACTTCATCTCTAGAGCTGGTGTGCGGTCTAGGGTTG
TGCATTTGGTTGGTTCTTTCGGCAAGAACGTGCGGGGTTCTATTAACCTGGTGACCGCCATCAAGGACAACTTCTCTAACGG
GCTTGTGTTCAAGGGCAACATTTTCGATATCAAGGCCAAAAAGACCCGTGAGAGCCTGGACAACTACCTGTCTATCTGCACC
ACACTTAGCCAGGCTCCTATCACCAAGCACGACAAGAACCAGATCCTGCGGTCACTTTTCGTTAGCGGTCCTCGGATTCAGT
ACGTGAGTTCTCAGTTTGGCTCTAGGCGGAACCGGATGTCTATTCTTCAAGAGGTTGTGGCTGACGACCCTACTCTTCATTG
GCCAGATCAGGATACCAGCCAGAAGCAGCTTGAGGATAAGTTCAGGGAACTCGCCCACAAAGAGTTGCCTTTCTTGACCGA

GAAGGTTTTCCACGACCACCTCGAGAAGATTGAGCAGCTTATGAAAGAAAACACCCACCTCGGTGGCAGGGATGTTGATGC
TTCTAAGACTCCTTACGTGCTGGCTAGGGCTAACGATATTGAGATCCACTGCTACGAGCTTTGGCGTGAGTATGATGAAGATG
AGGACGAGGCTTACCAGGCATACTGTTCTGAAGTTGAGGCCGCTATGGACCAAGAAAAGCTGAACGCTCTGATCGAGAGGT
ATCACGTTGACCCTAAGGCCAACTGGATCCAAATGCTTATGAACGGCGAGATCGAGACTGTCGAAGAGTTGAACAAGCTGG
ACAAGGGCTTCGAGTCTCACAGGTTGGCTCTTGTTGAGAGGATTCGTGTGGGTAAGCTTGGCATTCTTGGCAGCTATACAAA
GTGCCAGCAGCGAATCGAGGAACTTGATGGTGAGGGTAACAAGACCCATAGGTACACCGGTGAAGGCATTTGGAGGGGCTC
ATTCGATGATTCTGACGTGTGCATCGTTGTGCAGGACCTGAAAAAGACCAGAGAAAGCTACCTCAAGTGCGTCGTGTTCTCC
AAGGTTTCCGATTACAAGGTTCTGATGGGCCACCTCAAGACTTGGTGCAGGGAACACCATATCAGCAACGATGAGTTCCCG
ACTTGTACCCAGAAAGAACTCCTGTCTTACGGCGTGACCAAGTCCTCTGTGCTGCTCTACAAGATGAACGGCATGAAGATGC
TGCGGAACATGGAAAAGGGCATCCCACTTTACTGGAACCCGAGCCTTTCAACAAGGTCCCAGACCTACATTAACTGGCTGG
CTGTGGACATCACCGATCACTCACTTAGGCTTAGGAATAGGACCGTCGAAAACGGCAGGGTTGTGAACCAGACTATTATGGT
GGTGCCGCTGTACAAGACTGACGTGCAGATATTCAAGACCTCTCCGGTTGATCTCGAGCAGGATGTTCAGAACGATAGGCTG
AAGCTTCTGTCAGTGACTAAGGCTGGTGAGCTTAGATGGCTGCAGGACTGGATAATGTGGCGGTCATCTGCTGTGGACGACC
TCAACATTCTGAATCAGGTCAGGCGTAACAAGGCTGCTAGGGATCATTTCAACGCCAAGCCGGAATTCAAGAAGTGGATCA
AAGAGCTTTGGGACTACGCCCTGGATACCACTCTGATTAACAAGAAAGTGTTCATCACTACCCAGGGCAGCGAGTCTCAGTC
TACTGTTTCAAGCGGCGATTCCGATTCTGCTGTTGCTCCTCTTACTGATGAGGCCGTGGATGAGATCCATGACCTGCTTGACA
AAGAATTGGAGAAGGGGACCCTGAAGCAGATTATCCACGATGCTACCATTGACGCCCAGCTGGACATTCCTGCTATTGAGTC
TTTCCTGGCCGAAGAAATGGAAGTGTTCAAGTCCAGCCTCGCTAAGTCTCACCCTCTGCTTTTGAACTACGTGCGGTACATG
ATTCAAGAGATCGGGGTGACCAACTTCCGTAGCCTGATCGATAGCTTCAATCAGAAGGACCCGCTCAAGTCCGTGTCCTTGT
CTATCATCGATCTGAAAGAGGTTTTCAAGTTCGTCTACCAGGACATCAACGACGCCTACTTCGTTAAGCAAGAGGAAGATCA
CAAGTTCGACTTCTAG
